# Acquisition of cellular properties during alveolar formation requires differential activity and distribution of mitochondria

**DOI:** 10.1101/2021.04.09.439195

**Authors:** Kuan Zhang, Erica Yao, Julia Wong, Paul J. Wolters, Pao-Tien Chuang

## Abstract

Alveolar formation requires coordinated movement and interaction between alveolar epithelial cells, mesenchymal myofibroblasts and endothelial cells/pericytes to produce secondary septa. These processes rely on the acquisition of distinct cellular properties to enable ligand secretion for cell-cell signaling and initiate morphogenesis through cell migration and cell shape change. In this study, we showed that mitochondrial activity and distribution play a key role in bestowing cellular functions on both alveolar epithelial cells and mesenchymal myofibroblasts for generating secondary septa to form alveoli. These results suggest that mitochondrial function is tightly regulated to empower cellular machineries in a spatially specific manner. Indeed, such regulation via mitochondria is required for secretion of platelet-derived growth factor from alveolar epithelial cells to influence myofibroblast proliferation and migration. Moreover, mitochondrial function enables myofibroblast migration during alveolar formation. Together, these findings yield novel mechanistic insights into how mitochondria regulate pivotal steps of alveologenesis. They highlight selective utilization of energy and diverse energy demands in different cellular processes during development. Our work serves as a paradigm for studying how mitochondria control tissue patterning.

## Introduction

Production of alveoli during development and following lung injury is essential for lung function (1-3). Defective alveologenesis underlies bronchopulmonary dysplasia (BPD) (4) and ongoing destruction of alveoli is characteristic of chronic obstructive lung disease (COPD) (5). COPD is a major cause of morbidity and mortality globally (6, 7). During alveolar formation, alveolar epithelial cells (type I [AT1] and type II [AT2] cells), myofibroblasts and endothelial cells/pericytes undergo coordinated morphogenetic movement to generate secondary septa within saccules. As a result, secondary septa comprise of a layer of alveolar epithelial cells that ensheathes a core of myofibroblasts and endothelial cells/pericytes. Secondary septa formation (or secondary septation) is the most important step during alveolar formation. Platelet-derived growth factor (PDGF) produced by alveolar epithelial cells is a key player in controlling myofibroblast proliferation and migration during alveologenesis (8, 9). In response to PDGF signaling, myofibroblasts migrate to the prospective site of secondary septation and secrete elastin. Myofibroblasts and endothelial cells/pericytes are subsequently incorporated with alveolar epithelial cells to form secondary septa. All of these principal components play a key role in driving secondary septa formation (2). Generation of alveoli increases the surface area and efficiency of gas exchange, enabling high activity in terrestrial environments. Despite the progress that has been made, our mechanistic understanding of alveologenesis remains incomplete.

Mitochondrial activity is essential for every biological process and mitochondria provide a major source of ATP production through oxidative phosphorylation (OXPHOS) (10, 11). Unexpectedly, we have limited mechanistic insight into how mitochondria control cellular processes *in vivo*. In this regard, little is known about if and how mitochondria regulate alveolar formation. Many genetic and molecular tools have been developed in mice to study mitochondrial function. They offer a unique opportunity to address the central question of how mitochondria control alveologenesis at the molecular level.

Mitochondria exhibit dynamic distribution within individual cells. This process is mediated by the cytoskeletal elements that include microtubules, F-acitn and intermediate filaments. For instance, *Miro1* (*Mitochondrial Rho GTPase 1*), which is also called *Rhot1* (*ras homolog family member 1*), encodes an atypical Ras GTPase and plays an essential role in mitochondrial transport (12). Miro1 associates with the Milton adaptor (TRAK1/2) and motor proteins (kinesin and dynein), and tethers the adaptor/motor complex to mitochondria. This machinery facilitates transport of mitochondria via microtubules within mammalian cells. Whether regulated mitochondrial distribution is essential for lung cell function during alveologenesis is unknown.

In this study, we have demonstrated a central role of mitochondrial activity and distribution in conferring cellular properties to alveolar epithelial cells and myofibroblasts during alveolar formation. In particular, PDGF ligand secretion from alveolar epithelial cells and motility of myofibroblasts depend on regulated activity and distribution of mitochondria. Moreover, loss of mitochondrial function does not have a uniform effect on cellular processes, indicating diverse energy demands *in vivo*. We also reveal regulation of mitochondrial function by the mTOR complex 1 (13, 14) during alveolar formation and establish a connection between mitochondria and COPD/emphysema. Taken together, these findings provide new insight into how different cell types meet unique energy demands to engender distinct cellular properties during alveolar formation.

## Results

### Mitochondria display dynamic subcellular distribution in alveolar epithelial cells and mesenchymal myofibroblasts during alveolar formation

To uncover the functional role of mitochondria during alveologenesis, we first examined the distribution of mitochondria in murine lung cells involved in alveolar formation. We used antibodies against mitochondrial components to visualize the distribution of mitochondria in lung epithelial cells and myofibroblasts. For instance, we performed immunostaining on lung sections derived from *Sox9^Cre/+^; ROSA26^mTmG/+^* mice with anti-MPC1 (mitochondrial pyruvate carrier 1) and anti-MTCO1 (mitochondrially encoded cytochrome C oxidase I) (15). In particular, anti-MPC1 serves as a general marker for mitochondria. Lung epithelial cells were labeled by GFP produced from the *ROSA26^mTmG^* reporter (16) due to selective Cre expression in SOX9^+^ epithelial cells (17). We found that mitochondria were widely distributed in alveolar epithelial cells (distinguished by T1α and SPC for AT1 and AT2 cells, respectively) and myofibroblasts (marked by PDGFRA, PDGF receptor a, and smooth muscle actin, SMA) (18, 19) (Figure 1A). This is consistent with an essential role of mitochondrial activity in proper functioning of lung cells. In addition, we observed an uneven subcellular distribution of mitochondria (Figure 1A, 1B). For instance, mitochondria were concentrated in areas adjacent to the trans-Golgi network (TGN38^+^) in alveolar epithelial cells where proteins were sorted to reach their destinations through vesicles and in areas that surrounded SMA in myofibroblasts (Figure 1A, 1B). This finding suggests that localized mitochondrial distribution is required for cellular functions in mammalian lungs.

**Figure 1.**
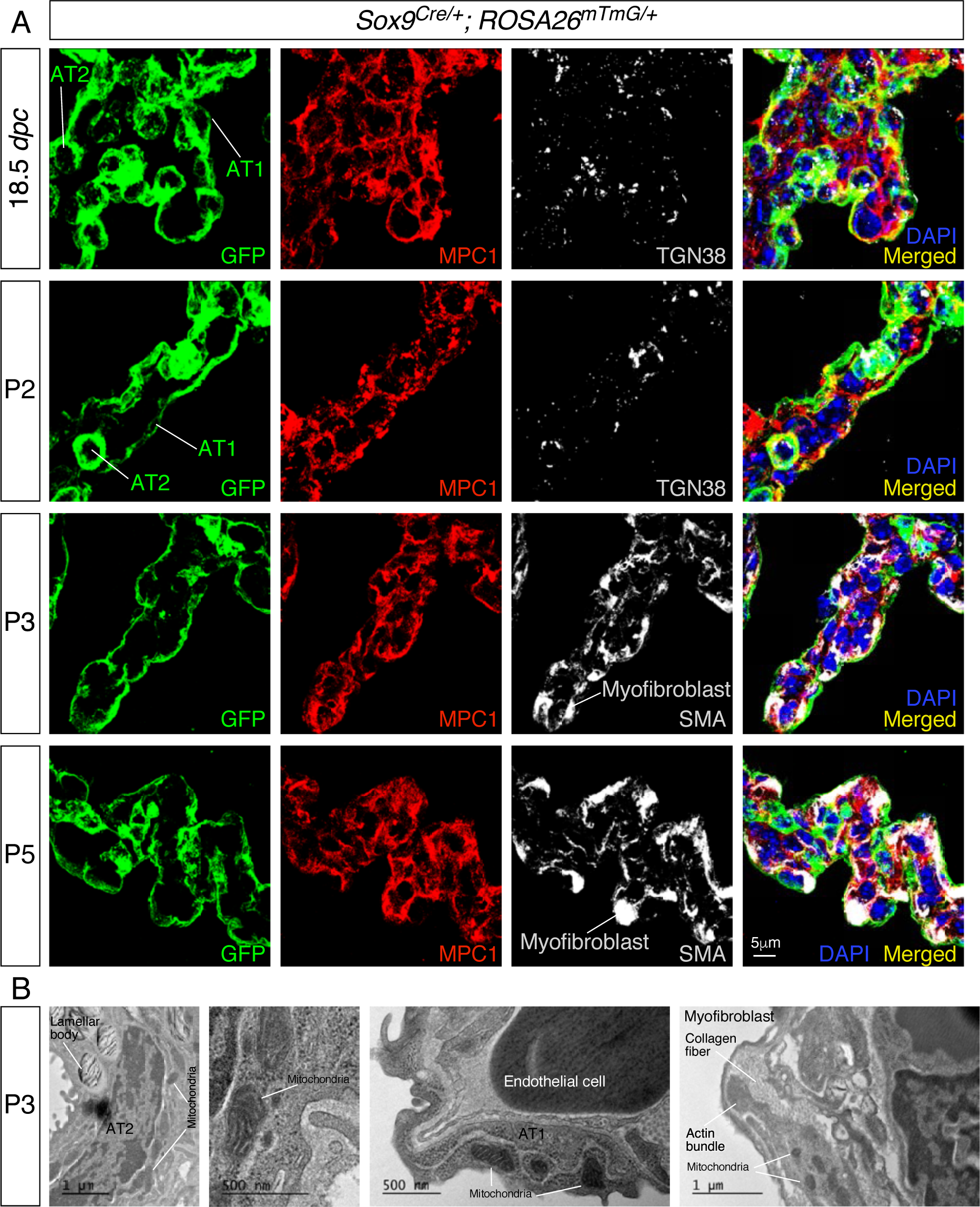
Mitochondria display subcellular concentration in alveolar epithelial cells and mesenchymal myofibroblasts of mouse lungs. (A) Immunostaining of lung sections collected from *Sox9^Cre/+^; ROSA26^mTmG/+^* mice at 18.5 *days post coitus* (*dpc*) and different postnatal (P) stages as indicated. The GFP signal identified alveolar epithelial cells while myofibroblasts were characterized by SMA expression. Moreover, mitochondria were labeled by MPC1; the trans-Golgi network was visualized by TGN38. Enhanced MPC1 signal was distributed non-uniformly in both alveolar epithelial cells and myofibroblasts. (B) Transmission electron micrographs of lungs collected from wild-type mice at P3. Prominent features in a given lung cell type include lamellar bodies in alveolar type II (AT2) cells, elongated cell membrane in alveolar type I (AT1) cells, and actin bundles and collagen fibers in myofibroblasts.

### Postnatal removal of mitochondrial activity in the lung leads to defective alveologenesis

We first tested if mitochondrial activity is required for alveologenesis by inactivating *Tfam* (transcription factor A, mitochondria), which encodes a master regulator of mitochondrial transcription (20), in the mouse lung after birth. We produced *CAGG^CreER/+^; ROSA26^mTmG/+^* (control) and *Tfam^f/f^; CAGG^CreER/+^; ROSA26^mTmG/+^* mice. Tamoxifen was administered to neonatal mice to activate CreER and lungs were collected at postnatal (P) day 10 (Figure 2A). CreER expression under the *CAG* promoter/enhancer (21) was ubiquitous in lung cells, including NKX2.1^+^ epithelial cells and PDGFRA^+^ fibroblasts/myofibroblasts, and converted a floxed allele of *Tfam* (*Tfam^f^*) (22) into a null allele (Figure 2B). We noticed that multiple regions in the lungs of mutant mice displayed alveolar defects concomitant with an increased mean linear intercept (MLI), a measure of air space size (23, 24) (Figure 2C, 2D). Alveolar defects were associated with disorganized SMA (Figure 2E). We anticipated that *Tfam* removal led to shutdown of mitochondrial transcription and loss of mitochondrial activity. Indeed, the relative ratio of mitochondrial DNA (mtDNA) (25), 16S rRNA and mtND1 (mitochondrially encoded NADH dehydrogenase 1), to nuclear DNA (nDNA), Hk2 (hexokinase 2), was reduced in *Tfam*-deficient lungs compared to controls (Figure 2F). Loss of *Tfam* was accompanied by diminished immunoreactivity of MTCO1, the expression of which was controlled by *Tfam*. (Figure 2G). Together, these results indicate that mitochondrial activity is required for alveolar formation.

**Figure 2.**
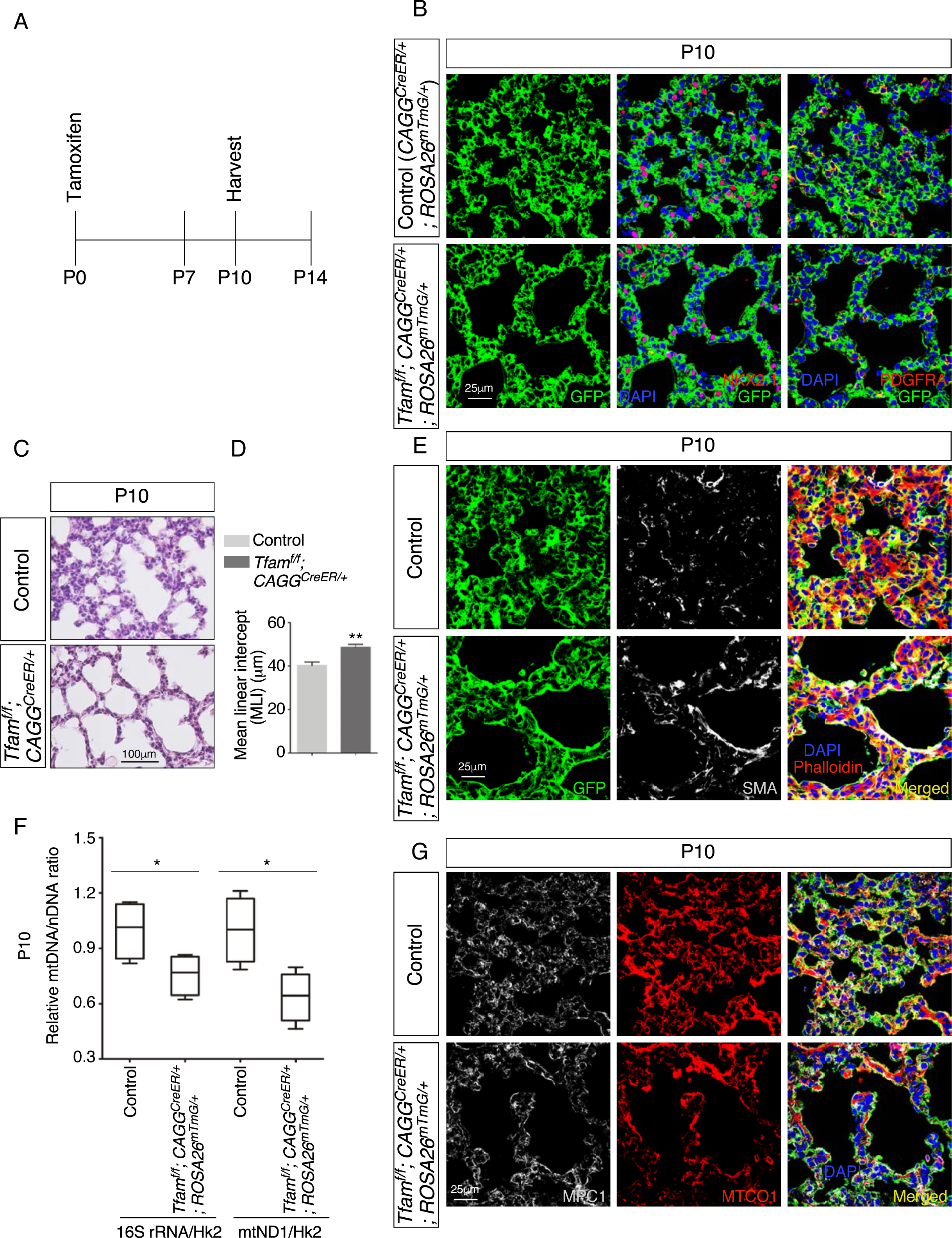
Global inactivation of *Tfam* in postnatal mice results in alveolar defects. (A) Schematic diagram of the time course of postnatal (P) administration of tamoxifen and harvest of mouse lungs. (B) Immunostaining of lungs collected from *CAGG^CreER/+^; ROSA26^mTmG/+^* (control) and *Tfam^f/f^*; *CAGG^CreER/+^; ROSA26^mTmG/+^* mice at P10 that had received tamoxifen at P0. The GFP signal represents sites of induced CreER activity. Nuclear NKX2.1 staining marked all lung epithelial cells while PDGFRA immunoreactivity labeled mesenchymal myofibroblasts. (C) Hematoxylin and eosin-stained lung sections of control and *Tfam^f/f^*; *CAGG^CreER/+^; ROSA26^mTmG/+^* mice at P10. Histological analysis revealed the presence of enlarged saccules and retarded development of secondary septa in the mutant lungs. (D) Measurement of the mean linear intercept (MLI) in control and *Tfam^f/f^*; *CAGG^CreER/+^; ROSA26^mTmG/+^* lungs at P10 (n = 4 for each group). The MLI was increased in *Tfam*-deficient lungs. (E) Immunostaining of lung sections collected from control and *Tfam^f/f^*; *CAGG^CreER/+^; ROSA26^mTmG/+^* mice at P10. SMA expression was characteristic of myofibroblasts and phalloidin stained the actin filaments. (F) Quantification of the relative ratio of mitochondrial DNA (mtDNA), 16S rRNA and mtND1 (mitochondrially encoded NADH dehydrogenase 1), to nuclear DNA (nDNA), Hk2 (hexokinase 2), in lysates derived from control and *Tfam^f/f^*; *CAGG^CreER/+^; ROSA26^mTmG/+^* lungs (n = 4 for each group). (G) Immunostaining of lung sections collected from control and *Tfam^f/f^*; *CAGG^CreER/+^; ROSA26^mTmG/+^* mice at P10. MPC1 antibodies marked mitochondria; MTCO1 antibodies detected cytochrome c oxidase, the expression of which was controlled by *Tfam*. All values are mean SEM. (*) p<0.05; (**) p<0.01 (unpaired Student’s *t*-test).

### Selective loss of mitochondrial activity in the lung epithelium disrupts alveologenesis

To investigate the function of mitochondrial activity in distinct compartments, we selectively eliminated mitochondrial activity in either the lung epithelium or mesenchyme. We produced control and *Tfam^f/f^; Sox9^Cre/+^* mice to establish a platform for mechanistic studies on mitochondrial activity in lung epithelial cells during alveolar formation. The *Sox9^Cre^* mouse line (17) is highly efficient in removing sequences flanked by loxP sites in the distal lung epithelium. *Sox9-Cre* is active at or later than 11.5 *days post coitus* (*dpc*) and converted *Tfam^f^* into a null allele and disrupted mitochondrial activity (Figure 3, figure supplement 1A). *Tfam^f/f^; Sox9^Cre/+^* mice were born at the expected Mendelian frequency and could not be distinguished from their wild-type littermates by their outer appearance or activity at birth. Moreover, histological analysis revealed no difference between control and mutant lungs prior to P5, confirming that branching morphogenesis and saccule formation were unaffected by inactivating *Tfam* in SOX9^+^ cells (Figure 3, figure supplement 2A). In addition, proliferation and differentiation of alveolar type I and type II cells proceeded normally in *Tfam^f/f^; Sox9^Cre/+^* lungs (Figure 3, figure supplement 2B). These results highlight a difference in dependence on mitochondrial activity in distinct cellular processes during development. To uncover the cellular processes that are highly dependent on mitochondrial activity, we investigated alveolar formation in control and *Tfam^f/f^; Sox9^Cre/+^* lungs.

**Figure 3.**
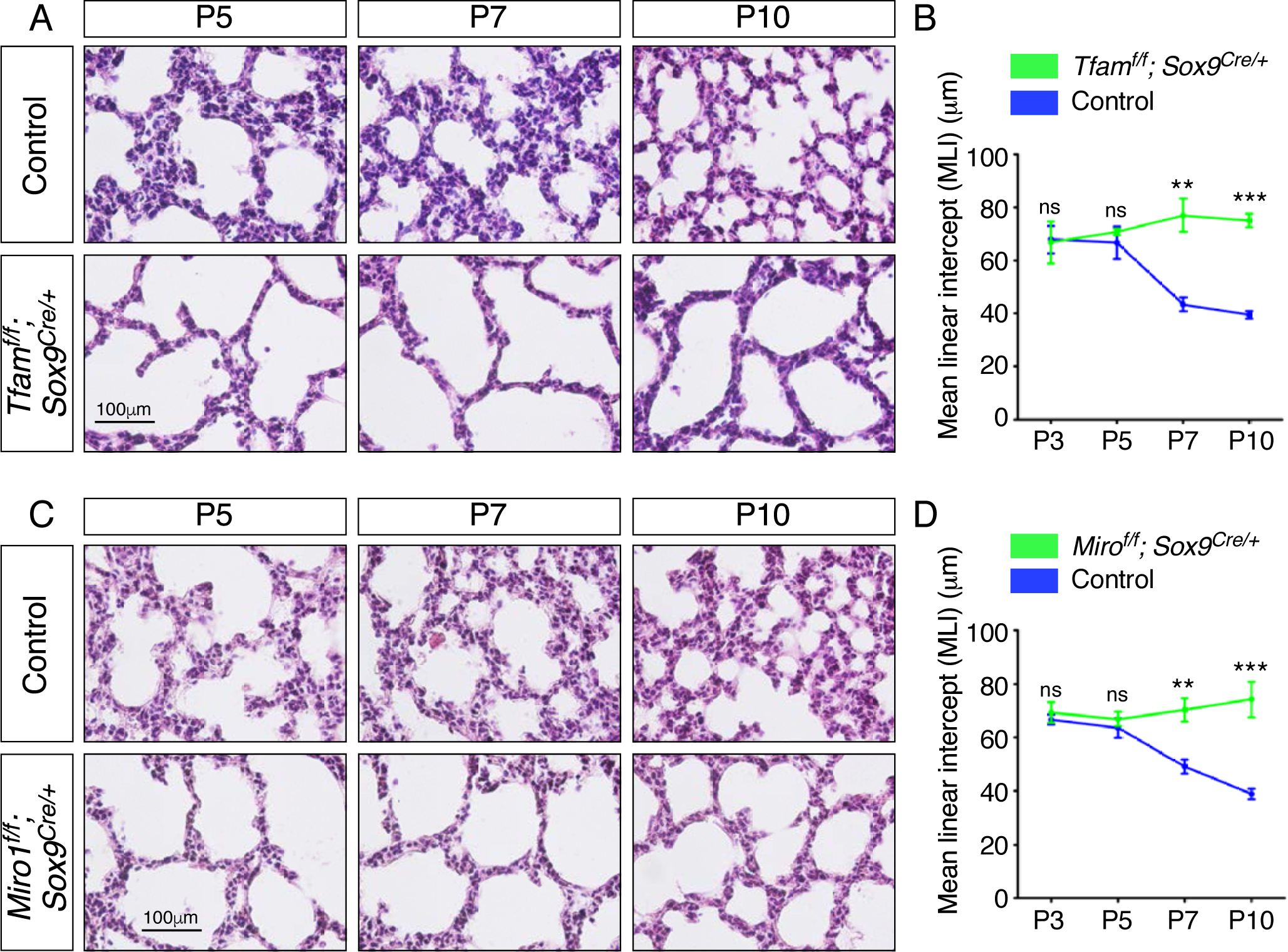
Elimination of *Tfam* or *Miro1* in the epithelium of mouse lungs disrupts alveolar formation. (A) Hematoxylin and eosin-stained lung sections of control and *Tfam^f/f^*; *Sox9^Cre/+^* mice at different postnatal (P) stages as indicated. Histological analysis revealed the presence of enlarged saccules and failure in secondary septation in the mutant lungs. (B) Measurement of the MLI in control and *Tfam^f/f^*; *Sox9^Cre/+^* lungs at P3–P10 (n = 3 for each group). The MLI was increased in *Tfam*-deficient lungs. (C) Hematoxylin and eosin-stained lung sections of control and *Miro1^f/f^*; *Sox9^Cre/+^* mice at different postnatal stages as indicated. Histological analysis detected enlarged saccules and lack of secondary septa in the mutant lungs. (D) Measurement of the MLI in control and *Miro^f/f^*; *Sox9^Cre/+^* lungs at P3–P10 (n = 3 for each group). The MLI was increased in *Miro1*-deficient lungs. All values are mean SEM. (**) p<0.01; (***) p<0.001; ns, not significant (unpaired Student’s *t*-test).

After P5, *Tfam^f/f^; Sox9^Cre/+^* mice could be discerned by their slightly reduced body size in comparison with the littermate controls. Histological analysis of *Tfam^f/f^; Sox9^Cre/+^* lungs at various postnatal stages revealed defects in secondary septa formation with an increased MLI (Figure 3A, 3B). Most *Tfam^f/f^; Sox9^Cre/+^* mice survived beyond P30, permitting analysis of the progression of their alveolar defects. These results reveal a critical role of mitochondrial activity in lung epithelial cells during alveologenesis.

### Disruption of mitochondrial distribution in the lung epithelium disturbs alveologenesis

As described above, mitochondria display dynamic distribution in lung cells, raising the possibility that proper subcellular distribution of mitochondria is vital for cellular function during alveolar formation. To test this hypothesis, we generated control and *Miro1^f/f^; Sox9^Cre/+^* mice. *Sox9^Cre^* converted a floxed allele of *Miro1* (*Miro^f^*) (26) to a null allele in SOX9^+^ alveolar epithelial cells. Loss of *Miro1* is expected to perturb normal subcellular distribution of mitochondria. *Miro1^f/f^; Sox9^Cre/+^* mice were born at the expected Mendelian frequency and cannot be distinguished from their wild-type littermates by their outer appearance or activity at birth. Similarly, no difference between control and mutant lungs prior to P5 was detected by histological analysis (Figure 3, figure supplement 2C, 2D). After P5, *Miro1^f/f^; Sox9^Cre/+^* mice displayed defects in secondary septation with an increased MLI. The alveolar phenotypes could first appear anywhere between P5 and P12 (Figure 3C, 3D). These results support the notion that localized mitochondrial distribution plays a functional role in alveolar formation. We noticed that the alveolar defects in *Miro1^f/f^; Sox9^Cre/+^* lungs were less severe than those in *Tfam^f/f^; Sox9^Cre/+^* lungs. This is likely due to the fact that only the distribution and not the activity of mitochondria was perturbed in *Miro1^f/f^; Sox9^Cre/+^* lungs.

### PDGF signal reception is perturbed and the number of mesenchymal myofibroblasts is reduced in the absence of proper mitochondrial activity or distribution in the lung epithelium

We examined various lung cell types in *Tfam^f/f^; Sox9^Cre/+^* lungs to explore the molecular basis of their alveolar phenotypes. Interestingly, the number of myofibroblasts marked by PDGFRA was reduced in the absence of epithelial *Tfam* (Figure 4A, 4D). Likewise, we found that the number of myofibroblasts was reduced in *Miro1^f/f^; Sox9^Cre/+^* lungs where epithelial *Miro1* was lost (Figure 4G, 4I). A diminished population of myofibroblasts in *Tfam^f/f^; Sox9^Cre/+^* and *Miro1^f/f^; Sox9^Cre/+^* lungs prompted us to investigate whether PDGF signaling was disrupted. Phosphorylation of PDGFRA (p-PDGFRA), indicative of PDGF signaling, was reduced in *Tfam^f/f^; Sox9^Cre/+^* or *Miro1^f/f^; Sox9^Cre/+^* lungs (Figure 4A, 4G). This observation suggests that PDGF signal reception by myofibroblasts was impaired in *Tfam^f/f^; Sox9^Cre/+^* or *Miro1^f/f^; Sox9^Cre/+^* lungs. Defective PDGF signal reception in myofibroblasts could be due to lack of PDGF production, trafficking or release.

**Figure 4.**
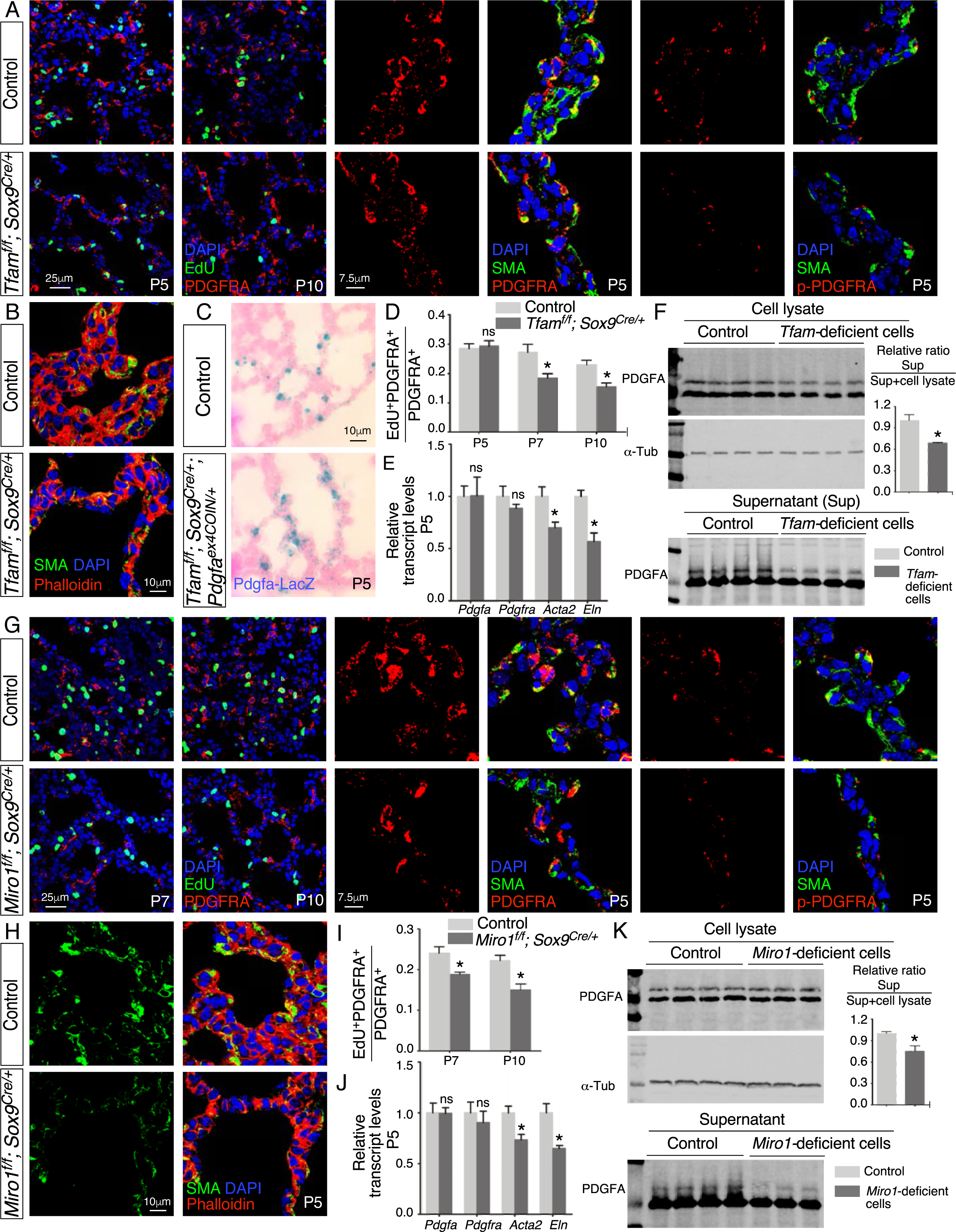
Loss of epithelial *Tfam* or *Miro1* compromises PDGF release. (A) Immunostaining of lungs collected from control and *Tfam^f/f^; Sox9^Cre/+^* mice at postnatal (P) day 5 or 10, some which were injected with EdU as indicated. (B) Immunostaining of lungs collected from control and *Tfam^f/f^; Sox9^Cre/+^* mice at P5. (C) LacZ-staining (blue) of lung sections collected from *Sox9^Cre/+^; Pdgfa^ex4COIN/+^* (control) and *Tfam^f/f^; Sox9^Cre/+^; Pdgfa^ex4COIN/+^* mice. The slides were counterstained with eosin (red). No difference in the intensity of LacZ staining in the lung was noted between these two mouse lines. (D) Quantification of myofibroblast proliferation in control and *Tfam^f/f^; Sox9^Cre/+^* lungs at P5, P7 and P10 (n = 3 for each group). The rate of myofibroblast proliferation was calculated as the ratio of the number of EdU^+^ myofibroblasts (EdU^+^PDGFRA^+^) to the number of myofibroblasts (PDGFRA^+^). The percentage of proliferating myofibroblasts was reduced in *Tfam^f/f^; Sox9^Cre/+^* compared to controls at P7 and P10. (E) qPCR analysis of gene expression in control and *Tfam^f/f^; Sox9^Cre/+^* lungs at P5 (n = 3 for each group). While no difference in expression levels was noted for *Pdgfa* and *Pdgfra* between control and *Tfam^f/f^; Sox9^Cre/+^* lungs, the expression levels of *Acta2* (SMA) and *Eln* (elastin) were significantly reduced in the absence of *Tfam*. (F) Western blot analysis of cell lysates and supernatants from control and *Tfam*-deficient cells (n = 4 for each group) lentivirally transduced with PDGFA-expressing constructs. The amount of PDGFA released into the media was reduced in *Tfam*-deficient cells compared to controls. α-Tubulin served as a loading control. (G) Immunostaining of lungs collected from control and *Miro1^f/f^; Sox9^Cre/+^* mice at P5, P7 or P10, some which were injected with EdU as indicated. (H) Immunostaining of lungs collected from control and *Miro1^f/f^; Sox9^Cre/+^* mice at P5. (I) Quantification of myofibroblast proliferation in control and *Miro1^f/f^; Sox9^Cre/+^* lungs at P7 and P10 (n = 3 for each group). The percentage of proliferating myofibroblasts was reduced in *Miro1^f/f^; Sox9^Cre/+^* compared to controls at P7 and P10. (J) qPCR analysis of gene expression in control and *Miro1^f/f^; Sox9^Cre/+^* lungs at P5 (n = 3 for each group). The expression levels of *Pdgfa* and *Pdgfra* were unaltered between control and *Miro1^f/f^; Sox9^Cre/+^* lungs; the expression levels of *Acta2* and *Eln* were significantly reduced in the absence of *Miro1*. (K) Western blot analysis of cell lysates and supernatants from control and *Miro1*-deficient cells (n = 4 for each group) lentivirally transduced with PDGFA-expressing constructs. The amount of PDGFA released into the media was reduced in *Miro1*-deficient cells compared to controls. α-Tubulin served as a loading control. All values are mean SEM. (*) p<0.05; ns, not significant (unpaired Student’s *t*-test).

We found that production of the PDGF ligand (PDGFA) in alveolar epithelial cells was unaffected in *Tfam^f/f^; Sox9^Cre/+^* or *Miro1^f/f^; Sox9^Cre/+^* lungs by qPCR analysis (Figure 4E, 4J). To substantiate this model, we utilized a PDGF reporter mouse line (*Pdgfa^ex4COIN^*) (27) that faithfully recapitulates the spatial and temporal expression of *Pdgfa*. Of note, no reliable PDGF antibody is available to detect PDGF in lungs or other tissues (19, 27). We generated *Pdgfa^ex4COIN/+^; Sox9*^Cre/+^ (control) and *Tfam^f/f^; Pdgfa^ex4COIN/+^; Sox9^Cre/+^* mice. Cre recombinase activated *β-galactosidase* (*lacZ*) expression in *Pdgfa*-expressing cells from the *Pdgfa^ex4COIN^* allele. We found that *LacZ* expression in *Pdgfa*-expressing cells (*i.e.*, alveolar epithelial cells) displayed a similar pattern and intensity between control and *Tfam*-deficient lungs (Figure 4C). Together, these results pointed to disrupted PDGF trafficking or release. This defect would subsequently disturb signal reception in mesenchymal myofibroblasts of *Tfam^f/f^; Sox9^Cre/+^* and *Miro1^f/f^; Sox9^Cre/+^* lungs.

### PDGF secretion from lung cells is diminished without proper mitochondrial activity or distribution

Our model posits that secretion of PDGF ligand from *Tfam*- and *Miro1*-deficient alveolar epithelial cells is compromised. To test this idea, we derived *Tfam*- and *Miro1*-deficient cells from *Tfam^f/f^; Pdgfra^Cre/+^* and *Miro1^f/f^; Pdgfra^Cre/+^* lungs (see below), respectively. We transduced control and *Tfam*- and *Miro1*-deficient cells with lentiviruses that produced epitope-tagged PDGF. Using this assay, we determined the amount of PDGF released from control and *Tfam*- and *Miro1*-deficient cells (Figure 4F, 4K). PDGF levels in the conditioned media derived from *Tfam*- or *Miro1*-deficient cells were reduced compared to controls (Figure 4F, 4K). These findings support a model in which loss of mitochondrial activity or distribution results in a failure of vesicular transport and PDGF release from alveolar epithelial cells. We surmise that these defects are in part due to an incapacitated actomyosin cytoskeleton caused by reduced mitochondrial function.

### Selective loss of mitochondrial activity or distribution in myofibroblasts compromises alveologenesis

We then investigated the functional requirement of mitochondria in the lung mesenchyme. To this end, we produced control and *Tfam^f/f^; Pdgfra^Cre/+^* (28) mice for *Tfam* inactivation in lung fibroblasts/myofibroblasts. Cre expression in PDGFRA^+^ myofibroblasts eliminated *Tfam* and mitochondrial activity. Indeed, the expression of MTCO1, a transcriptional target of *Tfam*, was reduced in lung fibroblasts/myofibroblasts derived from *Tfam^f/f^; Pdgfra^Cre/+^* mice (Figure 3, figure supplement 1B). Prior to P3, *Tfam^f/f^; Pdgfra^Cre/+^* mice could not be distinguished from their wild-type littermates by appearance, activity, morphological and immunohistochemical analysis (Figure 5, figure supplemental 1A, 1B). This suggests that loss of *Tfam* in the lung mesenchyme did not affect branching morphogenesis or saccule formation (29). This permitted us to assess the contribution of mesenchymal mitochondria to alveolar formation. Histological analysis of *Tfam^f/f^; Pdgfra^Cre/+^* lungs at various postnatal stages prior to P5 revealed defective secondary septa formation with an increased MLI (Figure 5A, 5B). Most *Tfam^f/f^; Pdgfra^Cre/+^* mice died before P30.

**Figure 5.**
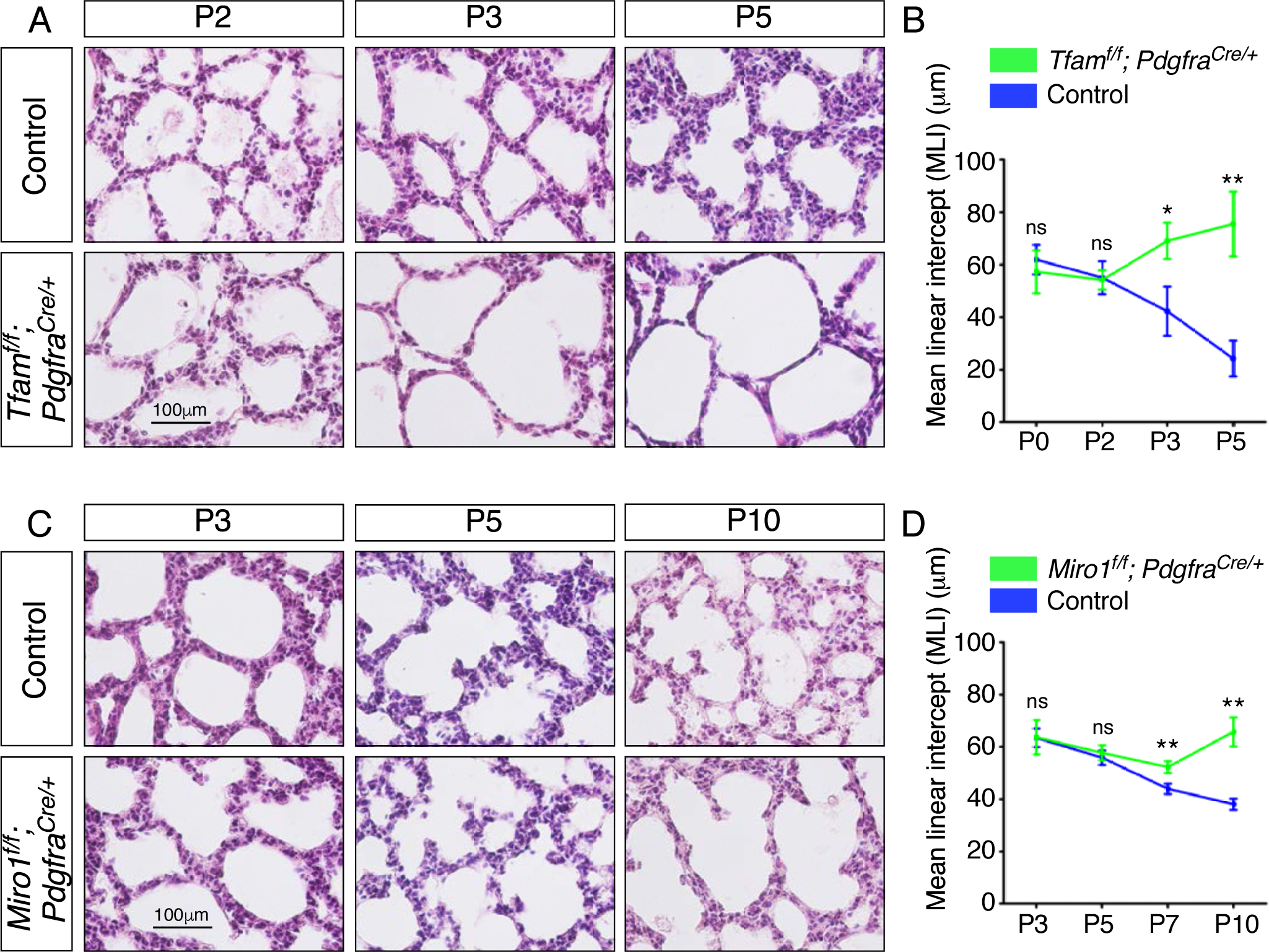
Elimination of *Tfam* or *Miro1* in mouse lung fibroblasts/myofibroblasts impairs alveolar formation. (A) Hematoxylin and eosin-stained lung sections of control and *Tfam^f/f^*; *Pdgfra^Cre/+^* mice at different postnatal (P) stages as indicated. Histological analysis revealed the presence of enlarged saccules and defective development of secondary septa in the mutant lungs. (B) Measurement of the MLI in control and *Tfam^f/f^*; *Pdgfra^Cre/+^* lungs at P0–P5 (n = 3 for each group). The MLI was increased in *Tfam*-deficient lungs. (C) Hematoxylin and eosin-stained lung sections of control and *Miro1^f/f^*; *Pdgfra^Cre/+^* mice at different postnatal stages as indicated. Histological analysis detected enlarged saccules and lack of secondary septa in the mutant lungs. (D) Measurement of the MLI in control and *Miro^f/f^*; *Pdgfra^Cre/+^* lungs at P3–P10 (n = 3 for each group). The MLI was increased in *Miro1*-deficient lungs. All values are mean SEM. (*) p<0.05; (**) p<0.01; ns, not significant (unpaired Student’s *t*-test).

We also generated control and *Tfam^f/f^; Dermo1^Cre/+^* mice, in which *Tfam* was eliminated by mesenchymal *Dermo1^Cre^* (30). *Tfam^f/f^; Dermo1^Cre/+^* mice exhibited alveolar defects (Figure 5, figure supplemental 2A, 2B) similar to those in *Tfam^f/f^; Pdgfra^Cre/+^* mice, further supporting a central role of mitochondrial activity in the lung mesenchyme during alveologenesis.

Of note, we bred *Lrpprc^f/f^; Pdgfra^Cre/+^* mice as an alternative means to disrupt mitochondrial activity. *Lrpprc* (*Leucine-rich PPR motif-containing*) (31) is required for mitochondrial translation. Similarly, *Pdgfra^Cre^* converted a floxed allele of *Lrpprc* (*Lrpprc^f^*) into a null allele. *Lrpprc* affects different aspects of mitochondrial activity and serves the purpose of confirming our findings using *Tfam*. We found that *Lrpprc^f/f^; Pdgfra^Cre/+^* mice developed alveolar defects (Figure 5, figure supplemental 2C, 2D), albeit the phenotypes were less severe than those in *Tfam^f/f^; Pdgfra^Cre/+^* mice. This was likely due to the presence of residual proteins after removal of *Lrpprc*. Together, these studies establish a crucial role of mitochondrial activity in myofibroblasts for alveologenesis.

We went on to determine whether proper subcellular distribution of mitochondria in myofibroblasts is necessary for their function during alveolar formation. We generated control and *Miro1^f/f^; Pdgfra^Cre/+^* mice (Figure 5, figure supplemental 1C, 1D). In this case, the subcellular distribution of mitochondria is expected to be perturbed in mesenchymal myofibroblasts as indicated by loss of proper MTCO1 distribution in myofibroblasts derived from lungs of *Miro1^f/f^; Pdgfra^Cre/+^* mice (Figure 3, figure supplement 1C). *Miro1^f/f^; Pdgfra^Cre/+^* mice exhibited alveolar defects, milder than those in *Tfam^f/f^; Pdgfra^Cre/+^* lungs (Figure 5C, 5D).

We discovered that myofibroblast proliferation was reduced in *Tfam^f/f^; Pdgfra^Cre/+^* or *Miro1^f/f^; Pdgfra^Cre/+^* lungs compared to controls (Figure 6A, 6B, 6E, 6F), suggesting defective PDGF signal reception. This may be related to a failure in PDGFR trafficking when mitochondrial function is impaired. As a result, the pool of PDGFRA^+^ myofibroblasts was decreased with a concomitant reduction in SMA (encoded by *Acta2*) and/or elastin (Figure 6C, 6G), contributing to alveolar defects. Taken together, these findings suggest that proper activity and distribution of mitochondria in mesenchymal myofibroblasts are critical to generating a sufficient number of myofibroblasts for secondary septation.

**Figure 6.**
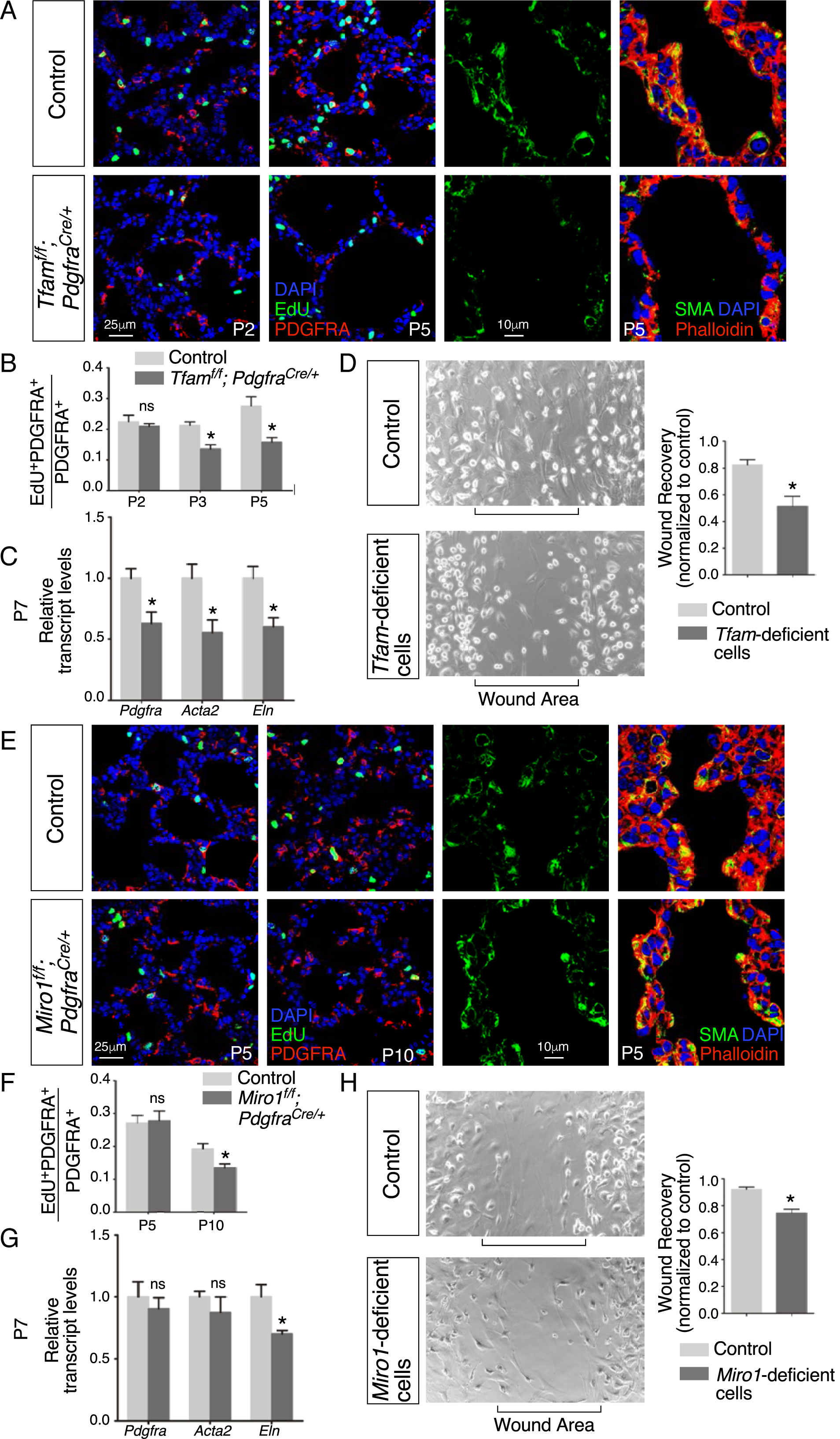
Loss of mesenchymal *Tfam* or *Miro1* compromises myofibroblast migration. (A) Immunostaining of lungs collected from control and *Tfam^f/f^; Pdgfra^Cre/+^* mice at postnatal (P) day 2 and 5, some of which injected with EdU as indicated. (B) Quantification of myofibroblast proliferation in control and *Tfam^f/f^; Pdgfra^Cre/+^* lungs at P2, P3 and P5 (n = 3 for each group). The rate of myofibroblast proliferation was calculated as the ratio of the number of EdU^+^ myofibroblasts (EdU^+^PDGFRA^+^) to the number of myofibroblasts (PDGFRA^+^). The percentage of proliferating myofibroblasts was reduced in *Tfam^f/f^; Pdgfra^Cre/+^* compared to controls at P3 and P5. (C) qPCR analysis of gene expression in control and *Tfam^f/f^; Pdgfra^Cre/+^* lungs at P3 (n = 3 for each group). The expression levels of *Pdgfra*, *Acta2* and *Eln* were significantly reduced in *Tfam^f/f^; Pdgfra^Cre/+^* lungs compared to controls. (D) Wound recovery assays to assess the migratory ability of myofibroblasts derived from control and *Tfam^f/f^; Pdgfra^Cre/+^* lungs (n = 3 for each group). Within 36–48 hr, the wound area has been populated by migrating myofibroblasts derived from control lungs. By contrast, fewer myofibroblasts from *Tfam^f/f^; Pdgfra^Cre/+^* lungs reached the wound area within the same time frame. Wound recovery by myofibroblasts from control and *Tfam^f/f^; Pdgfra^Cre/+^* lungs was quantified. (E) Immunostaining of lungs collected from control and *Miro1^f/f^; Pdgfra^Cre/+^* mice at P5 and P10, some of which were injected with EdU as indicated. (F) Quantification of myofibroblast proliferation in control and *Tfam^f/f^; Pdgfra^Cre/+^* lungs at P5 and P10 (n = 3 for each group). The percentage of proliferating myofibroblasts was decreased in *Miro1^f/f^; Pdgfra^Cre/+^* compared to controls at P10. (G) qPCR analysis of gene expression in control and *Miro1^f/f^; Pdgfra^Cre/+^* lungs at P7 (n = 3 for each group). The expression levels of *Eln* were significantly reduced in *Miro1^f/f^; Pdgfra^Cre/+^* lungs in comparison with controls. (H) Wound recovery assays to assess the migratory ability of myofibroblasts derived from control and *Miro1^f/f^; Pdgfra^Cre/+^* lungs (n = 3 for each group). Fewer myofibroblasts from *Miro1^f/f^; Pdgfra^Cre/+^* lungs reached the wound area within the same time frame compared to controls. Wound recovery by myofibroblasts from control and *Miro1^f/f^; Pdgfra^Cre/+^* lungs was quantified. All values are mean SEM. (*) p<0.05; ns, not significant (unpaired Student’s *t*-test).

### Migration of myofibroblasts is reduced without proper mitochondrial activity or distribution, which is associated with a disrupted cytoskeleton

We speculate that myofibroblasts deficient in *Tfam* are also defective in their ability to migrate to the prospective site of secondary septation. To test this idea, we isolated myofibroblasts from control and *Tfam^f/f^; Pdgfra^Cre/+^* lungs and seeded them onto the migration chamber for wound healing assays (32). The rate of myofibroblast migration into the cell-free area was measured. While control myofibroblasts occupied the cell-free area after 36–48 hr, only scant *Tfam*-deficient myofibroblasts were detected in the cell-free area (Figure 6D). This result indicates that mobility of myofibroblasts was compromised due to loss of mitochondrial activity in these cells. We conjecture that the migration defect was in part due to an incapacitated actomyosin cytoskeleton without mitochondrial activity. Consistent with this idea, organization of the cytoskeleton (labeled by phalloidin) was perturbed in *Tfam^f/f^; Pdgfra^Cre/+^* lungs in comparison with controls (Figure 6A).

Similarly, myofibroblasts derived from *Miro1^f/f^; Pdgfra^Cre/+^* lungs displayed a compromised response in wound healing assays compared to controls (Figure 6H). Together, these results affirm the role of mitochondrial activity and distribution in regulating myofibroblast migration during alveologenesis.

### The mTOR complex 1 regulates mitochondrial function and alveologenesis

We have discovered an essential role of mitochondria in controlling alveolar formation. To dissect the signaling cascade that regulates mitochondrial function during alveologenesis, we first investigated mTOR complex 1 (mTORC1), a known regulator of mitochondrial activity and biogenesis. We generated control, *Rptor^f/f^; Sox9^Cre/+^* and *Rptor^f/f^; Pdgfra^Cre/+^* mice. *Rptor* (*Raptor*, *rapamycin-sensitive regulatory associated protein of mTOR*) (33) encodes an essential component of mTORC1, which includes RPTOR, mTOR and several other proteins. *Rptor^f/f^; Sox9^Cre/+^* and *Rptor^f/f^; Pdgfra^Cre/+^* mice failed to survive to term, precluding the analysis of potential alveolar phenotypes.

To circumvent this problem, we produced control and *Rptor^f/f^; CAGG^CreER/+^* mice and administered tamoxifen postnatally (Figure 7A). Analysis of *Rptor^f/f^; CAGG^CreER/+^* mice at P10 revealed alveolar defects with an increased MLI (Figure 7B, 7C). Defective secondary septation in *Rptor^f/f^; CAGG^CreER/+^* lungs was associated with reduced SMA (Figure 7D). As expected, the protein levels of phosphorylated ribosomal protein S6 (p-RPS6), a downstream target of mTORC1, was decreased in lysates from *Rptor*-deficient lungs compared to controls (Figure 7E). The relative ratio of mtDNA to nDNA was reduced in *Rptor*-deficient lungs (Figure 7F). Moreover, immunoreactivity of both MPC1 and MTCO1 was diminished in *Rptor^f/f^; CAGG^CreER/+^* lungs compared to controls (Figure 7G). These findings show that elimination of *Rptor* in the lung resulted in loss of mitochondria. Impaired mitochondrial function then contributed to alveolar defects. Together, our results suggest that mTORC1 controls alveolar formation partly through its effects on mitochondrial function.

**Figure 7.**
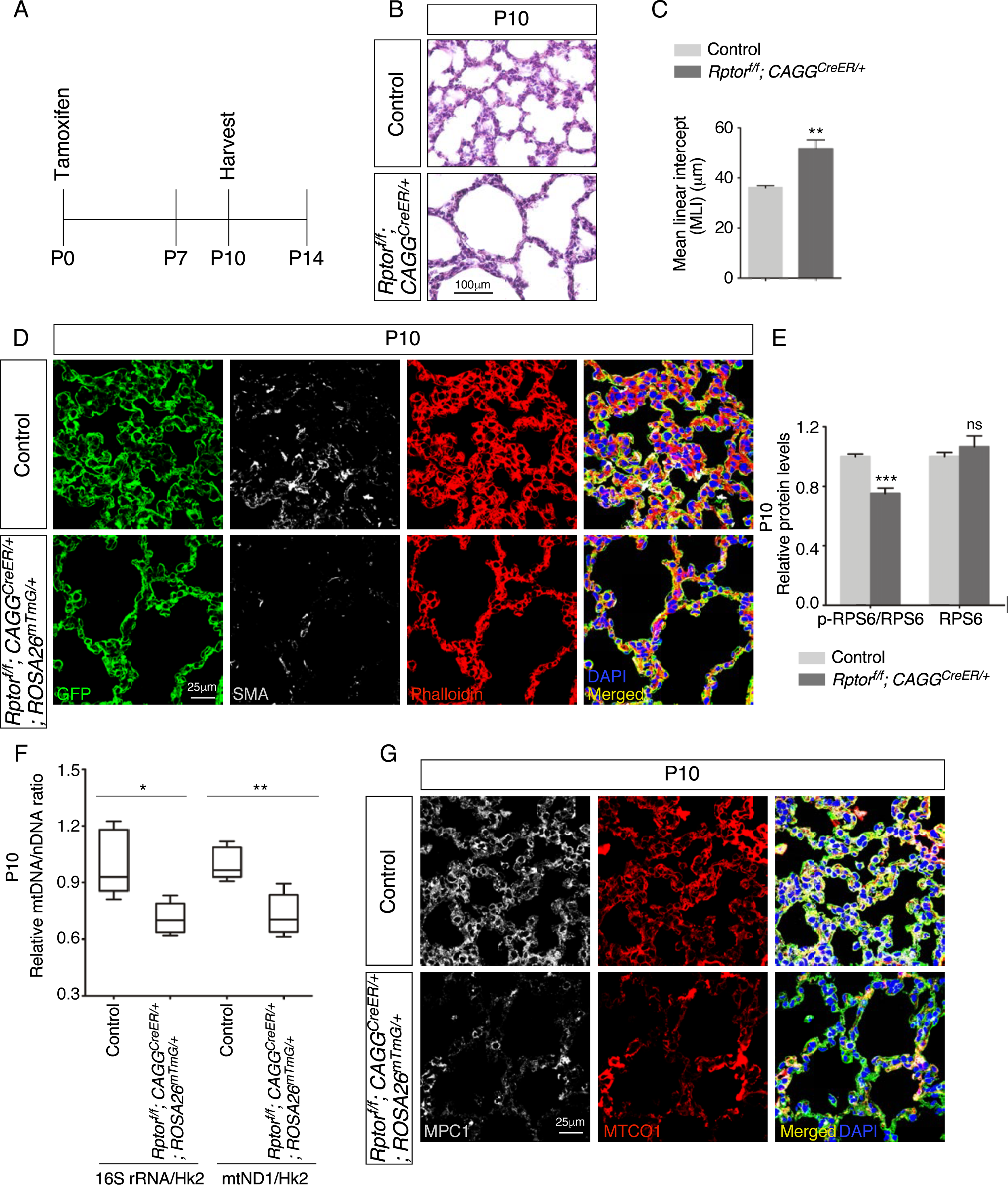
Postnatal inactivation of *Rptor* in mice results in alveolar defects. (A) Schematic diagram of the time course of postnatal (P) administration of tamoxifen and harvest of mouse lungs. (B) Hematoxylin and eosin-stained lung sections of control and *Rptor^f/f^*; *CAGG^CreER/+^* mice at postnatal (P) day 10. Histological analysis revealed the presence of enlarged saccules and retarded development of secondary septa in the mutant lungs. (C) Measurement of the mean linear intercept (MLI) in control and *Rptor^f/f^*; *CAGG^CreER/+^* lungs at P10 (n = 5 for each group). The MLI was increased in *Rptor*-deficient lungs. (D) Immunostaining of lung sections collected from control and *Rptor^f/f^*; *CAGG^CreER/+^; ROSA26^mTmG/+^* mice at P10. SMA expression was characteristic of myofibroblasts and phalloidin stained the actin filaments. (E). Quantification of the protein levels of ribosomal protein S6 (RPS6) and phosphorylated RPS6 (p-RPS6) in lung lysates derived from control and *Rptor^f/f^*; *CAGG^CreER/+^; ROSA26^mTmG/+^* lungs (n = 5 for each group). (F) Quantification of the relative ratio of mitochondrial DNA (mtDNA), 16S rRNA and mtND1 (mitochondrially encoded NADH dehydrogenase 1), to nuclear DNA (nDNA), Hk2 (hexokinase 2), in lysates derived from control and *Rptor^f/f^*; *CAGG^CreER/+^; ROSA26^mTmG/+^* lungs (n = 5 for each group). (G) Immunostaining of lung sections collected from control and *Rptor^f/f^*; *CAGG^CreER/+^; ROSA26^mTmG/+^* mice at P10. MPC1 antibodies marked mitochondria; MTCO1 antibodies detected cytochrome c oxidase, the expression of which was controlled by *Tfam*. All values are mean SEM. (*) p<0.05; (**) p<0.01; (***) p<0.001 (unpaired Student’s *t*-test).

### Mitochondrial copy number and TFAM protein levels are decreased in lungs from COPD/emphysema patients

To explore whether studies of mitochondrial function in alveolar formation in mice can shed new light on human diseases, we assessed the status of mitochondria in lung tissues of normal subjects and COPD/emphysema patients (Figure 8A). We found that the copy number of mitochondria (25) relative to nuclear DNA was significantly reduced in COPD/emphysema patients (Figure 8B). In addition, TFAM protein levels were reduced in lysates from emphysema lungs compared to normal lungs (Figure 8C). These results established a connection between mitochondrial function and pathogenesis of COPD/emphysema. Interestingly, a disorganized cytoskeleton was noted in COPD/emphysema lungs, in which actin bundles seen in normal alveoli were sparse (Figure 8D). We did not detect a difference in the protein levels of either ribosomal protein S6 (RPS6) or phosphorylated RPS6 (p-RPS6), a downstream target of mTORC1, in lysates from emphysema and normal lungs. However, we observed heterogeneous expression of p-RPS6 in emphysema lungs (Figure 8E). Whether regional reduction of p-RPS6 is correlated with disease progression needs future studies. Taken together, these findings complement our mouse work and lay the foundation for further investigation into the disease mechanisms of COPD/emphysema.

**Figure 8.**
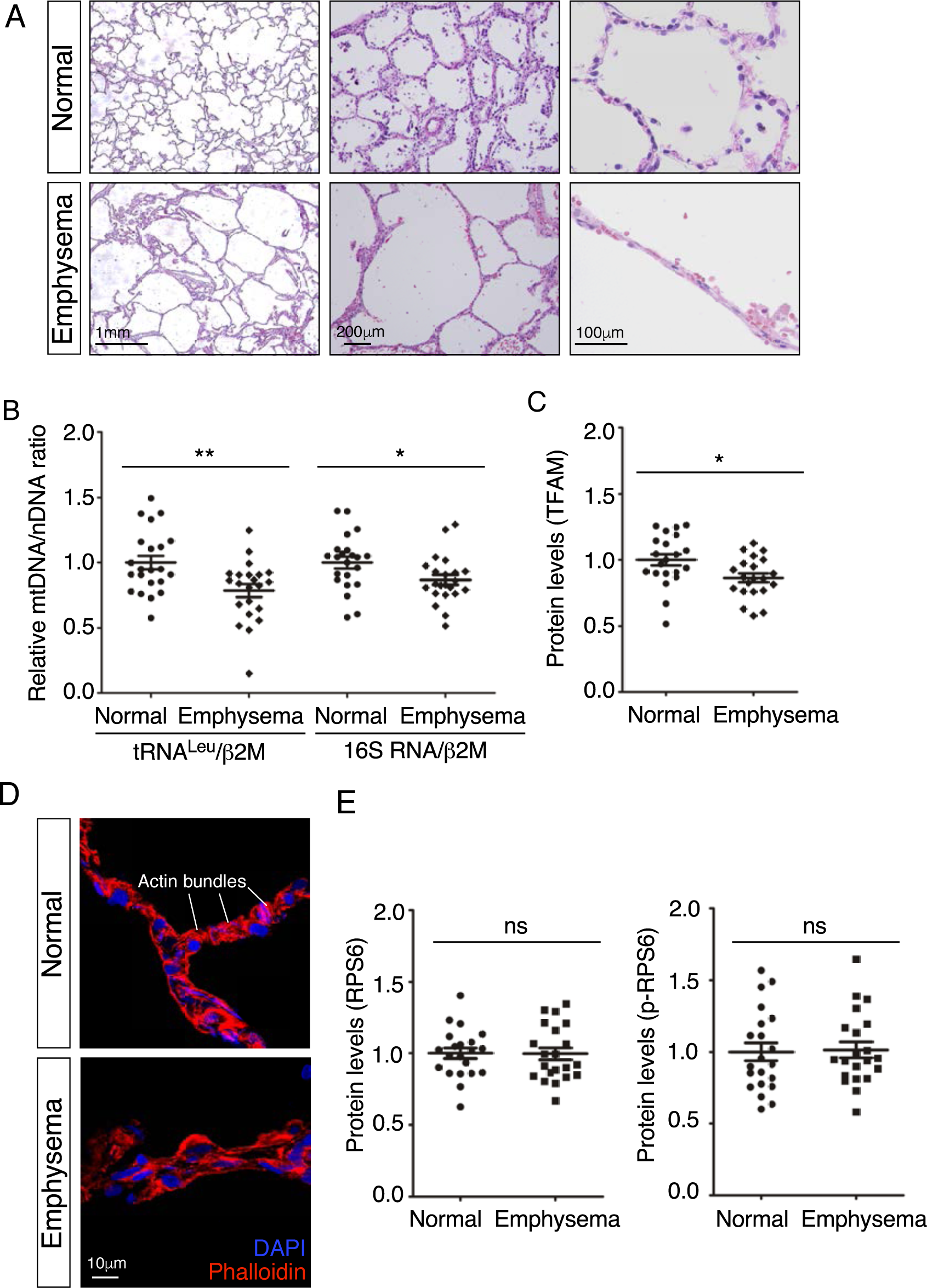
Lungs from emphysema patients exhibit a reduction in mitochondrial DNA and TFAM expression. (A) Hematoxylin and eosin-stained lung sections of normal and emphysema patients. Disruption of alveoli in emphysema patients resulted in enlarged airspace with thin primary septa. (B) qPCR analysis of mitochondrial DNA (mtDNA) to nuclear DNA (nDNA) in lung lysates derived from normal and emphysema patients (n = 22 for each group). mtDNA-encoded tRNA^Leu^ and 16S rRNA and nDNA-encoded β2μ (β2 microglobulin) were used in this study. The relative ratio of mtDNA to nDNA was calculated. (C) Quantification of TFAM protein levels in lung lysates derived from normal and emphysema patients (n = 21 for each group). (D) Immunostaining of lungs collected from control and emphysema patients. Phalloidin detected actin filaments. (E). Quantification of the protein levels of ribosomal protein S6 (RPS6) and phosphorylated RPS6 (p-RPS6) in lung lysates derived from normal and emphysema patients (n = 21 for each group). All values are mean SEM. (*) p<0.05; (**) p<0.01; ns, not significant (unpaired Student’s *t*-test).

## Discussion

Our studies have provided new insight into how mitochondrial function controls alveolar formation. We discovered a major role of mitochondria in conferring requisite cellular properties to both alveolar epithelial cells and myofibroblasts during alveologenesis. These findings define the molecular basis of the functional requirement of mitochondria in distinct compartments. They also add a new layer of complexity to the interactions between the major players of secondary septa. Moreover, our work establishes the foundation for investigating the interplay between mitochondria and signaling pathways in endowing cellular properties in alveologenesis during development and following injury. We expect that this framework will be applicable to other developmental systems (Figure 9).

**Figure 9.**
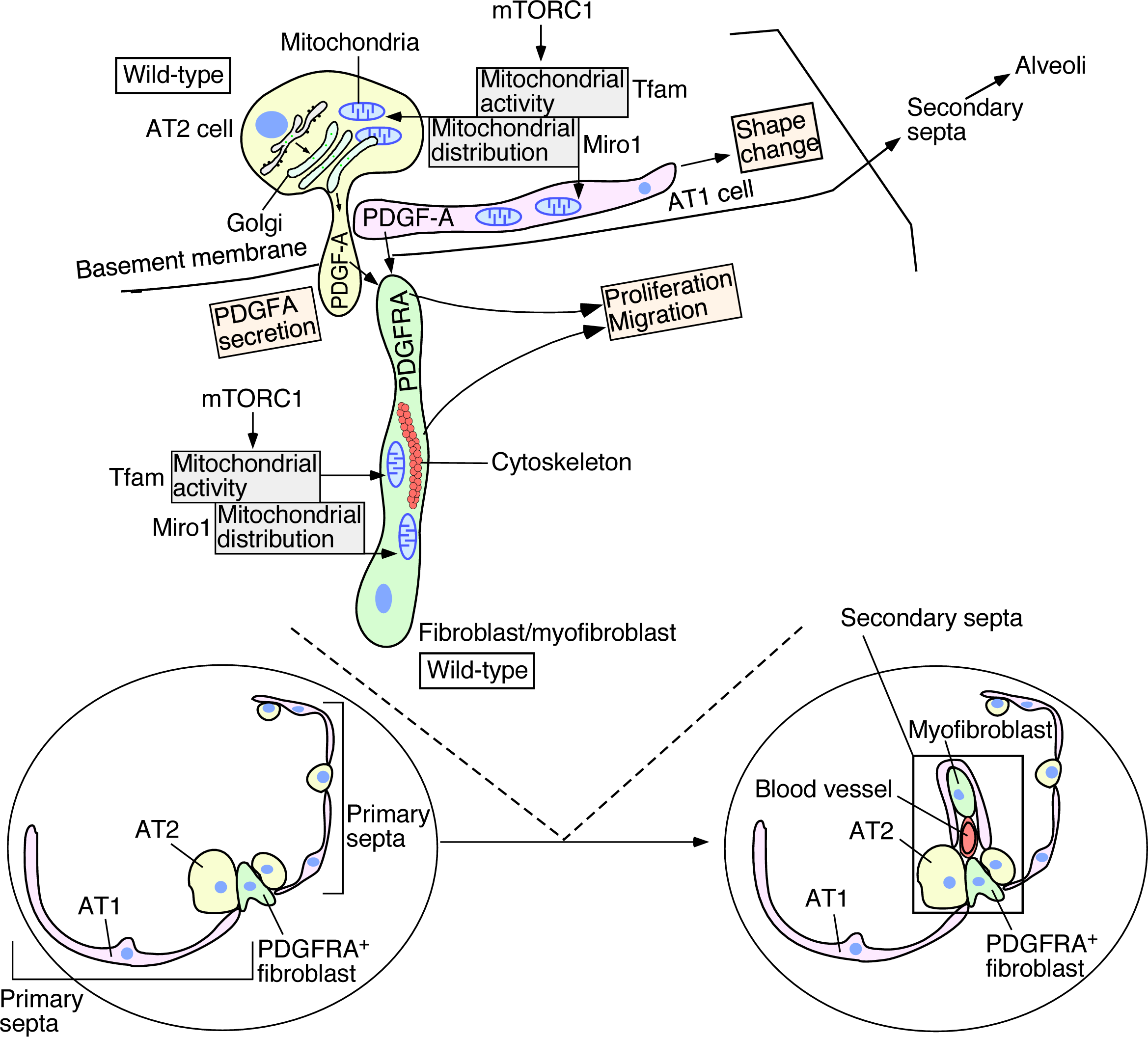
A model of regulating alveolar formation through mitochondrial activity and distribution. The main players during alveolar formation are shown in the schematic diagram. Mitochondria display dynamic subcellular distribution in in alveolar epithelial cells and mesenchymal fibroblasts/myofibroblasts. *Tfam* controls mitochondrial activity while *Miro1* regulates mitochondrial distribution. Mitochondrial activity and distribution in alveolar epithelial cells (type I [AT1] and type II [AT2]) contribute to secretion of the PDGF-A ligand. Reception of PDGF-A by mesenchymal myofibroblasts is critical to myofibroblast proliferation and migration, a key step in secondary septation. Similarly, mitochondrial activity and distribution in myofibroblasts are also required for myofibroblast contraction and migration, likely through powering the cytoskeleton. The mTORC1 pathway plays a central role in controlling mitochondrial function during alveolar formation. Specification of alveolar epithelial cells and myofibroblasts was unaffected in mutant mouse lungs in which mitochondrial activity and distribution were perturbed. We propose that essential cellular processes have differential requirements of mitochondrial activity and distribution. Investigating how mitochondria control signaling pathways and cellular processes *in vivo* provide a new way to functionally define signaling pathways and cellular processes. We surmise that the regulatory circuitry mediated by mitochondria activity and distribution is also deployed during alveolar repair following lung injury.

Removal of mitochondria either globally or in distinct compartments in the lung results in alveolar defects. We found that not all cellular processes are perturbed to the same extent when mitochondrial activity or distribution is perturbed. For instance, while alveolar development is disrupted, saccule formation and cell type specification are unaffected. These observations highlight the differential requirement of mitochondrial function in a given cellular process. Events that necessitate a higher demand of energy will be the first to exhibit phenotypes once mitochondrial function is compromised. It implies that the severity of phenotypes resulting from a reduction in mitochondrial function could be used to characterize the energy demand of a particular cellular process in development. This information cannot be obtained from cell-based studies.

Our work has focused on PDGF ligand secretion. While other pathways have not been interrogated, different pathways may exhibit a varying degree of dependence on mitochondrial function. Similarly, this could provide a new way to functionally categorize signaling pathways on the basis of their energy demand *in vivo*. Such insight will provide a new blueprint of how cells dispense its energy source in tissues.

Secretion of the PDGF ligand from alveolar epithelial cells is impaired due to mitochondrial dysfunction in either activity or distribution. We surmise that the actomyosin cytoskeleton likely underlies this defect. However, it is also possible that the energy produced by mitochondria is required in many key steps of vesicular transport and membrane fusion. Additional insight would come from a careful assessment of the dependence of each process on mitochondrial function. Similarly, failure to power the actomyosin cytoskeleton in myofibroblasts due to disruption of mitochondrial activity or distribution could cause the migratory defect. ATP produced by mitochondria is known for the assembly of the actomyosin cytoskeleton. By contrast, how regulation of mitochondrial distribution is superimposed upon the formation of the actomyosin cytoskeleton is less understood at the molecular level. This scenario is further complicated by the observation that F-actin and intermediate filaments also play a key role in mitochondrial dynamics and functions (34). Technical advances that enable live imaging (35) of mitochondria (36) and the cytoskeleton would provide new tools to address this important issue.

We have uncovered the role of mitochondria in alveolar epithelial cells and myofibroblasts during alveolar formation. Whether the function of endothelial cells/pericytes also relies on mitochondrial activity and/or distribution in this process is unknown (37, 38). This would require future studies using an approach similar to that employed in this work. Again, the dependence of cell types, cellular processes and signaling pathways on mitochondrial function can only be revealed through studies *in vivo*.

We envision that a complex regulatory network must be in place to regulate mitochondrial number and distribution in distinct cellular processes. Our work shows that mTOR complex 1 is a key player in controlling mitochondrial function during alveolar formation. This is consistent with the role of mTORC1 in regulating mitochondrial function in cell-based assays. Identifying additional components in the signaling network would reveal the key hubs in the signaling network that control mitochondrial function during alveologenesis.

Mitochondrial copy number and TFAM expression levels are reduced in the lungs of COPD/emphysema patients. However, it is unclear if mitochondrial activity and distribution contribute to the pathogenesis of COPD/emphysema. This would rely on the analysis of lungs at different stages of disease progression. It also necessitates new approaches that enable a mechanistic correlation between mitochondrial function and disease progression. For instance, analysis of the transcriptome and proteome of various lung cell types at single cell levels could reveal alterations in a subset of cells that herald the process of emphysematous changes.

Taken together, our work has yielded new molecular insight into the old question of energy utilization *in vivo*. In particular, energy production by mitochondria is channeled in a spatially specific manner to enable cellular processes in distinct cell types. Diverse cell types and cellular processes have a unique energy demand, which is likely executed by a signaling network. These investigations form the basis of additional studies to explore this new concept in alveolar formation and repair and in other processes *in vivo*.

## Materials and methods

### Animal husbandry

Mouse strains used in this study are listed in the Key Resources table. All mouse experiments described in this study were performed according to the protocol approved by the Institutional Animal Care and Use Committee (IACUC) of the University of California, San Francisco (UCSF).

### Tamoxifen administration

Tamoxifen was prepared by dissolving in corn oil to a concentration of 50 mg/ml (32). For postnatal (P) injection, the tamoxifen stock was diluted 1/10 in corn oil to make a final concentration of 5 mg/ml. 50 μL of tamoxifen was delivered through oral gavage or direct injection into the stomach of neonatal mice.

### Measurement of mean linear intercept (MLI)

Measurement of MLI was performed as previously reported (32). Briefly, 15 fields without visible blood vessels or airways from three different histological sections per animal were captured at 20x magnification using a Nikon Eclipse E1000 microscope. A grid with horizontal and vertical lines (10 each) was superimposed on the images using ImageJ. The mean linear intercept (Lm) was calculated as: Lm = L/N, where L is the total length of horizontal plus vertical lines, and N is the total number of the intercepts.

### Whole lung imaging, histology and immunohistochemistry

To image the whole lungs that carried GFP or RFP, dissected mouse lungs at the indicated stages were placed under a Nikon Eclipse E1000 microscope equipped with a SPOT 2.3 CCD camera.

Mouse lungs at indicated time points were collected and fixed with 4% paraformaldehyde (PFA) in PBS on ice for 1 hr. The tissues were embedded in paraffin wax or OCT, and sectioned at 7 μm. For histological analysis of lung sections, hematoxylin and eosin (H&E) staining was performed as previously described (32, 39). Images were taken using a SPOT 2.3 CCD camera connected to a Nikon Eclipse E1000 microscope.

To detect Pdgfa in lung cells, lungs from *Sox9^Cre/+^; Pdgfa^ex4COIN/+^* (control) and *Tfam^f/f^; Sox9^Cre/+^; Pdgfa^ex4COIN/+^* mice at P3 were dissected and fixed in 4% PFA on ice for 1 hr. Lungs were washed in 0.02% NP40 in PBS for 2 hr, then placed in X-gal staining solution (5 mM K_3_Fe(CN)_6_, 5 mM K_4_Fe(CN)_6_, 2 mM MgCl_2_, 0.01% sodium deoxycholate, 0.02% NP-40, 1 mg/ml X-gal) for 72 hr at 37°C. The stained lungs were paraffin embedded and sectioned. Images were taken using a SPOT 2.3 CCD camera connected to a Nikon Eclipse E1000 microscope.

Immunofluorescence was performed as previously described (39). Antibodies used in this study are listed in the Key Resources table. The primary antibodies used for wax sections were: chicken anti-GFP (1:200, abcam, Cat# ab13970), rabbit anti-NKX2.1 (1:100, Epitomics, Cat# 2044–1), goat anti-CC10 (1:200, Santa Cruz Biotechnology, Cat# sc-9773), mouse anti-acetylated tubulin (1:200, Sigma-Aldrich, Cat# T6793), rabbit anti-prosurfactant protein C (proSP-C) (1:200, MilliporeSigma, Cat# AB3786), hamster anti-T1α (1:200, Developmental Studies Hybridoma Bank, Cat# 8.1.1), mouse anti-HOPX (1:100, Santa Cruz Biotechnology, Cat# sc-398703). The primary antibodies used for frozen sections were: rabbit anti-MPC1 (1:100, Millipore/Sigma, Cat# HPA045119), rat anti-E-cadherin (1:200, Invitrogen, Cat# 13– 1900), mouse anti-MTCO1 (1:100, abcam, Cat# ab14705), chicken anti-GFP (1:300, abcam, Cat# ab13970), mouse anti-ACTA2 (1:200, Thermo Scientific Lab Vision, Cat# MS-113-P0), rat anti-PECAM-1 (CD31) (1:150, Santa Cruz Biotechnology, Cat# sc-18916), rabbit anti-PDGFRA (1:150, Cell Signaling Technology, Cat# 3164), rabbit anti-phospho-PDGFRA (Tyr754) (1:100, Cell Signaling Technology, Cat# 2992), mouse anti-S6 Ribosomal Protein (1:100, Cell Signaling Technology, Cat# 2317), rabbit anti-Phospho-S6 Ribosomal Protein (Ser235/236) (1:100, Cell Signaling Technology, Cat# 4856). Secondary antibodies and conjugates used were: donkey anti-rabbit Alexa Fluor 488 or 594 (1:1000, Life Technologies), donkey anti-chicken Alexa Fluor 488 or 647 (1:1000, Life Technologies), donkey anti-mouse Alexa Fluor 488 or 594 (1:1000, Life Technologies), and donkey anti-rat Alexa Fluor 594 (1:1000, Life Technologies). The biotinylated secondary antibodies used were: goat anti-hamster (1:1000, Jackson ImmunoResearch Laboratories), donkey anti-rabbit (1:1000, Jackson ImmunoResearch Laboratories), donkey anti-rat (1:1000, Jackson ImmunoResearch Laboratories) and horse anti-mouse (1:1000, Jackson ImmunoResearch Laboratories). The signal was detected using streptavidin-conjugated Alexa Fluor 488, 594, or 647 (1:1000, Life Technologies) or HRP-conjugated streptavidin (1:1000, Perkin-Elmer) coupled with fluorogenic substrate Alexa Fluor 594 tyramide for 30 s (1:200, TSA kit; Perkin Elmer). F-actin was stained with rhodamine-conjugated phalloidin (1:200, Sigma) in PBS for 2 hr. Since the filamentous actin is sensitive to methanol, ethanol and high temperature, we only used OCT-embedded frozen sections for F-actin staining.

Confocal images were captured on a Leica SPE laser-scanning confocal microscope. Adjustment of red/green/blue/grey histograms and channel merges were performed using LAS AF Lite (Leica Microsystems).

### Myofibroblast proliferation assays

The rate of cell proliferation was determined through EdU incorporation as previously described (32). Mouse pups at the indicated time points were intraperitoneally injected with EdU/PBS solution for 1hr before dissection. The Click-iT EdU Alexa Fluor 488 Imaging Kit (Life Technologies) was used to quantify EdU incorporation. The sections were co-stained with antibody against PDGFRA. Cell proliferation rate was calculated as the ratio of (EdU^+^PDGFRA^+^ cells)/(PDGFRA^+^ cells).

### Myofibroblast migration assay in vitro

The migratory capacity of lung myofibroblasts was assessed by the Culture-Insert 2 Well system (ibidi) (32). Briefly, at postnatal (P) day 3, dissected lungs from control, *Tfam^f/f^; Pdgfra^Cre/+^*and *Miro1^f/f^; Pdgfra ^Cre/+^* mice were minced into small pieces and digested in solution (1.2 U/ml dispase, 0.5mg/ml collagenase B and 50 U/ml DNase I), rocking at 37°C for 2 hr to release single cells. After adding an equal volume of culture medium (DMEM with 10% FBS, 2x penicillin/streptomycin and 1x L-glutamine), the samples were filtered through 40 μm cell strainers and centrifuged at 600 g for 10 min. The dissociated cells were resuspended in 200 μl of culture medium and plated into wells (100 μl per well). The lung myofibroblasts were allowed to attach to the fibronectin–coated plates for 2-3 hr. Fresh culture medium was added and the attached myofibroblasts were cultured 2 or 3 days to reach 100% confluence. The confluent myofibroblasts were switched to starvation medium (DMEM with 0.5% FBS and 1x penicillin/streptomycin) for 16 hr before removal of the insert. Myofibroblasts migration was assessed 36-48 hr afterwards.

### Lentiviral production and transduction

Lentiviruses were produced in HEK293T cells maintained in DMEM containing 10% FBS, 1x penicillin/streptomycin and 1x L-glutamine (32). HEK293T cells were plated and transfected when they reached 70% confluence on the following day. 2 μg of pMD2.G, 2 μg of psPAX2, 4 μg of PDGFA-3xFLAG (in the modified pSECC lentiviral vector) and 50 μl of polyethylenimine (PEI) (1 μg/μl) were mixed in 1000 μl OPTI-MEM and added to a 10 cm dish. Incubation medium was replaced one day after transfection. 48 hr post-transfection, the viral supernatant was collected, filtered through 0.45 μm PVDF filter, then added to primary lung cells together with 8 μg/ml polybrene. 12 hr post-transduction, the medium was replaced with fresh culture medium.

### PDGFA secretion assay

PDGFA secretion assay was performed as previously described (32). In brief, PDGFA-3xFLAG was stably expressed in control and *Tfam*- and *Miro1*-deficient primary lung cells (derived from the lungs of control, *Tfam^f/f^; Pdgfra^Cre/+^* and *Miro1^f/f^; Pdgfra^Cre/+^* mice, respectively) that were plated onto 10 cm dishes. The culture media were replaced with OPTI-MEM supplemented with insulin, transferrin and selenium (ITS) once the cells reached 100% confluence. 24 hr post-incubation, the supernatants were mixed with 1x protein inhibitor cocktail, filtered through 0.45 μm filters, and centrifuged at high speed (>12000 rpm) for 15 min at 4°C. The filtrates were then concentrated in protein concentration columns (Millipore CENTRICON YM-10 Centrifugal Filter Unit 2 mL 10 kDa) through centrifugation at 2000 g for 1 hr at 4°C. Concentrated supernatants were mixed with immunoprecipitation (IP) buffer (50 mM Tris pH 7.4, 2 mM EDTA, 150 mM NaCl, 0.5% Triton X-100, 1x protein inhibitor cocktail) in a total volume of 500 μl. Meanwhile, cells seeded on the plate were harvested and lysed in IP buffer. Immunoprecipitation was performed using FLAG-M2 beads following the standard procedure. Western blotting was performed to detect PDGFA in the supernatants and lysates derived from control, *Tfam*-deficient, and *Miro1*-deficient primary lung cells.

### qPCR analysis

qPCR was performed as previously described (32). Briefly, the right cranial lobe was dissected from the mouse lungs of the indicated genotypes and time points, and homogenized in 1 ml TRIzol (Life Technologies). The homogenates were added to 200 μl chloroform, and then centrifuged for 15 min at 12000 rpm. The upper aqueous layer was collected and mixed with an equal volume of 70% ethanol. RNAs were extracted with the RNeasy Mini Kit (Qiagen) following the manufacturer’s instructions. The extracted RNAs were reverse-transcribed with the Maxima First Strand cDNA Synthesis Kit (Thermo Scientific). Quantitative PCR (qPCR) was performed on the Applied Biosystems QuantStudio^TM^ 5 Real-Time PCR System. Primers used for qPCR were: mouse *Pdgfa* forward, 5’-CTGGCTCGAAGTCAGATCCACA-3’; reverse, 5’-GACTTGTCTCCAAGGCATCCTC-3’, mouse *Pdgfra* forward, 5’-TGCAGTTGCCTTACGACTCCAGAT-3’; reverse, 5’-AGCCACCTTCATTACAGGTTGGGA-3’, mouse *Acta2* forward, 5’-ATGCAGAAGGAGATCACAGC-3’; reverse, 5’-GAAGGTAGACAGCGAAGCC-3’, mouse *Eln* forward, 5’-GCCAAAGCTGCCAAATACG-3’; reverse, 5’-CTCCAGCTCCAACACCATAG-3’, mouse *Gapdh* forward, 5’-AGGTTGTCTCCTGCGACTTCA-3’; reverse, 5’-CCAGGAAATGAGCTTGACAAAGTT-3’.

### Analysis of mtDNA/nDNA ratio

Lung samples from *Tfam^f/f^; CAGG^CreER/+^*, *Rptor^f/f^; CAGG^CreER/+^* mice and human patients were lysed in lysis buffer (100 mM Tris pH 7.5, 5 mM EDTA, 0.4% SDS, 200 mM NaCl, 50 μg /ml proteinase K), and incubated in a 55°C chamber overnight. The tissues were digested with 100 μg/ml RNase A at 37°C for 30 min to degrade the RNAs. The lysed samples were then added with equal volume of phenol/chloroform and centrifuged at 12,000 rpm for 10 min. Lung DNAs were concentrated by ethanol precipitation. For mouse lungs, the mitochondrial 16S rRNA or ND1 gene and the nuclear *Hk2* gene were used to calculate the relative ratio of mitochondrial (mt) to nuclear (n) DNA copy number. For human lungs, the mitochondrial tRNA^Leu(UUR)^ or 16S rRNA gene and the nuclear β2-microglobulin (*β2M*) gene were employed to determine the relative ratio of mtDNA to nDNA. Quantitative PCR (qPCR) was performed on the Applied Biosystems QuantStudio^TM^ 5 Real-Time PCR System. The primer pairs used for the indicated genes were: mouse 16S rRNA forward, 5’-CCGCAAGGGAAAGATGAAAGAC-3’; reverse, 5’-TCGTTTGGTTTCGGGGTTTC-3’, mouse mt-ND1 forward, 5’-CTAGCAGAAACAAACCGGGC-3’; reverse, 5’-CCGGCTGCGTATTCTACGTT-3’, mouse Hk2 forward, 5’-GCCAGCCTCTCCTGATTTTAGTGT-3’; reverse, 5’-GGGAACACAAAAGACCTCTTCTGG-3’, human tRNA^Leu(UUR)^ forward, 5’-CACCCAAGAACAGGGTTTGT-3’; reverse, 5’-TGGCCATGGGTATGTTGTTA-3’, human 16S rRNA forward, 5’-GCCTTCCCCCGTAAATGATA-3’; reverse, 5’-TTATGCGATTACCGGGCTCT-3’, human β-2-microglobulin *(β2M*) forward, 5’-TGCTGTCTCCATGTTTGATGTATCT-3’; reverse, 5’-TCTCTGCTCCCCACCTCTAAGT-3’.

### Human lung tissues

Human lung samples were processed as previously described (32). Briefly, lung tissues were obtained from severe Emphysema (Global Initiative for Chronic Obstructive Lung Disease Criteria, stages III or IV) at the time of lung transplantation. The donor control lung samples were indicated physiologically and pathologically normal (40). Written informed consent was obtained from all subjects and the study was approved by the University of California, San Francisco Institutional Review Board (IRB approval # 13-10738).

### Western blotting analysis of human lung tissues

Human lung tissues from normal subjects and emphysema patients were homogenized in RIPA buffer with 1x Protease Inhibitor Cocktail and 1x PMSF. The lysates were centrifuged with full speed for 15 min at 4°C and analyzed by Western blotting as previously described (39). The primary antibodies used were: rabbit anti-TFAM (1: 2000, Proteintech, Cat# 22586-1-AP), mouse anti-S6 Ribosomal Protein (1:2000, Cell Signaling Technology, Cat# 2317), rabbit anti-Phospho-S6 Ribosomal Protein (Ser235/236) (1:2000, Cell Signaling Technology, Cat# 4856), mouse anti-alpha-tubulin (1:3000, Developmental Studies Hybridoma Bank, Cat# 12G10).

### Statistical analysis

All the statistical comparisons between different groups were shown as mean value ± SEM. The P values were calculated by two-tailed Student’s *t*-tests and statistical significance was evaluated as (*) p<0.05; (**) p<0.01; (***) p<0.001. More than or equal to three biological repeats were performed, and the detailed biological replicates (n numbers) were indicated in the figure legends.

## Author contributions

KZ, EY, and PTC conceived the project and designed the research. KZ, EY, JW and PTC performed the experiments and analyzed the data. PW provided human lung tissues and analyzed the data. KZ, EY and PTC drafted the manuscript. All authors read and reviewed the manuscript.

## Acknowledgments

Some data for this study were acquired at the Nikon Imaging Center at CVRI. This work was supported by R01 HL142876 from the National Institutes of Health to P.T.C.

---

**Figure 3–figure supplement 1.**
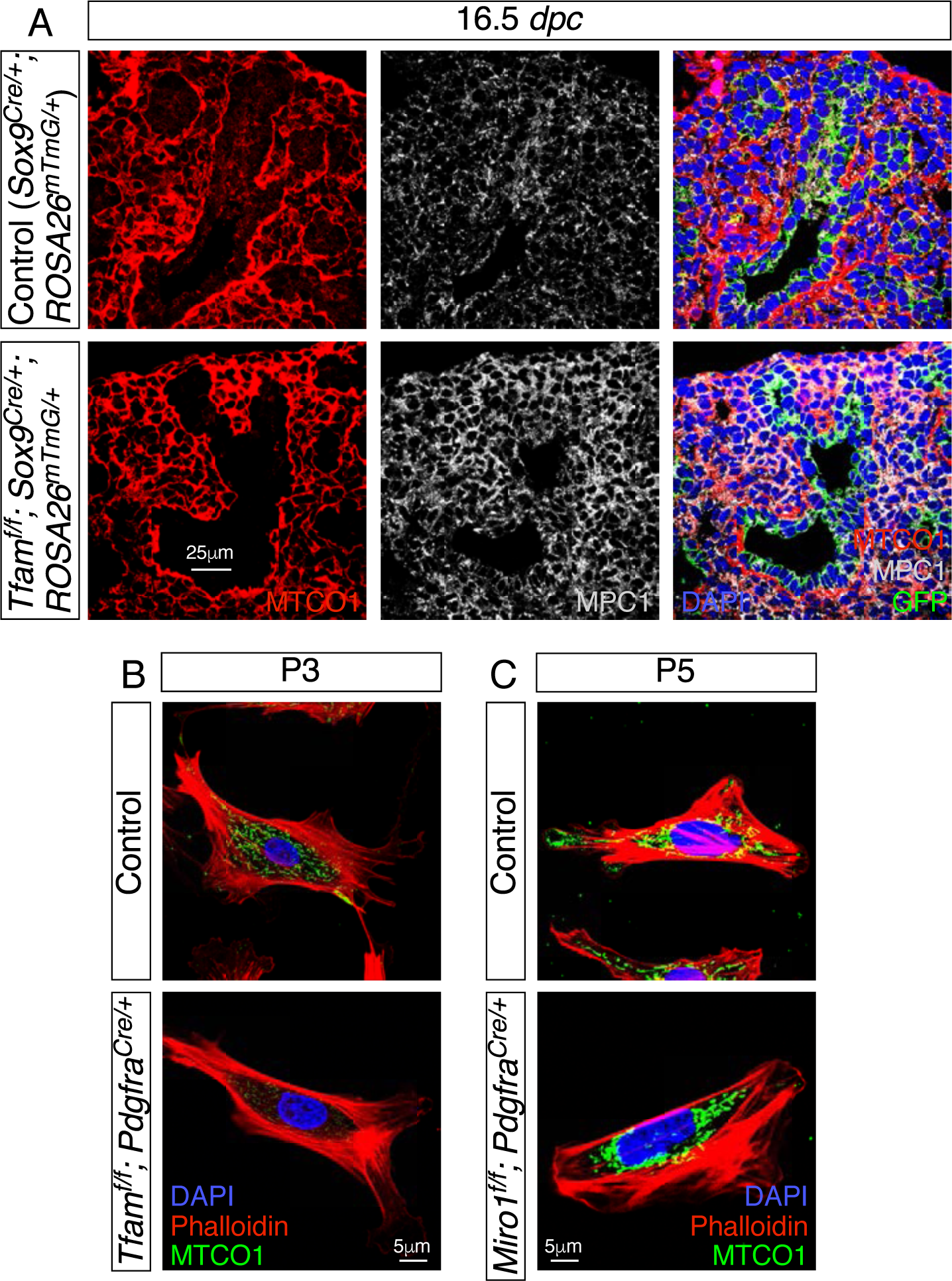
Loss *Tfam* or *Miro1* disrupts mitochondrial activity and distribution, respectively. (A) Immunostaining of lung sections collected from *Sox9^Cre/+^; ROSA26^mTmG/+^* (control) and *Tfam^f/f^; Sox9^Cre/+^; ROSA26^mTmG/+^* mice at 16.5 *days post coitus* (*dpc*). MPC1 antibodies marked mitochondria while MTCO1 antibodies detected cytochrome c oxidase, the expression of which was controlled by Tfam. (B) Immunostaining of fibroblasts derived from control or *Tfam^f/f^; Pdgfra^Cre/+^* lungs at postnatal (P) day 3. (C) Immunostaining of fibroblasts derived from control or *Miro1^f/f^; Pdgfra^Cre/+^* lungs at P5.

**Figure 3–figure supplement 2.**
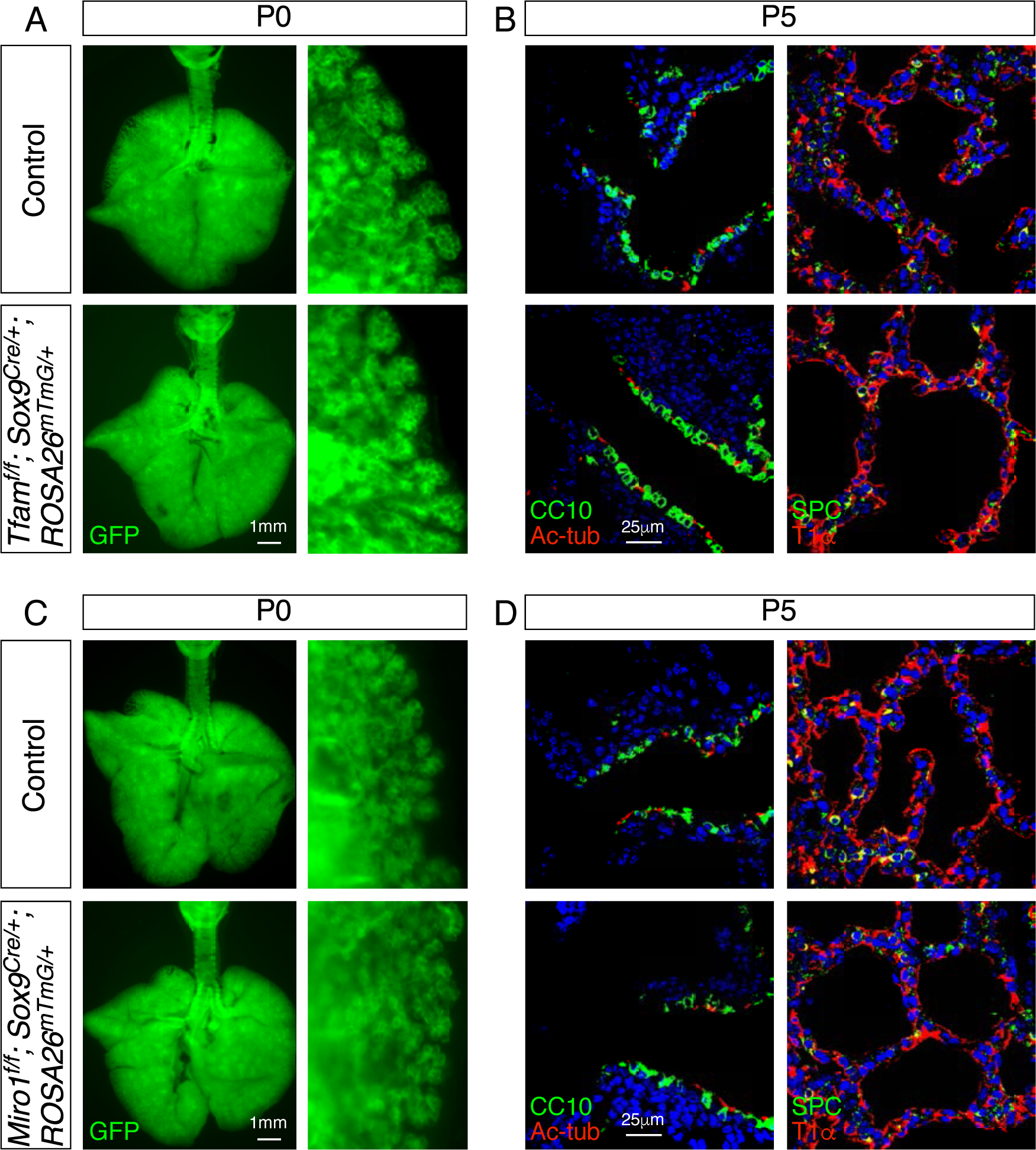
Elimination of *Tfam* or *Miro1* in the epithelium of mouse lungs does not perturb saccule formation or cell type specification. (A) Surface view of dissected lungs from *Sox9^Cre/+^; ROSA26^mTmG/+^* (control) and *Tfam^f/f^; Sox9^Cre/+^; ROSA26^mTmG/+^* mice at postnatal (P) day 0. No difference in saccule formation was noted between control and mutant lungs. (B) Immunostaining of lung sections from control and *Tfam^f/f^; Sox9^Cre/+^; ROSA26^mTmG/+^* mice at P5. Specification of lung cell types (e.g., club cell [CC10^+^], ciliated cell [Ac-tub^+^], AT1 cell [T1^+^], AT2 cell [SPC^+^]) were unaffected. (C) Surface view of dissected lungs from *Sox9^Cre/+^; ROSA26^mTmG/+^* (control) and *Miro1^f/f^; Sox9^Cre/+^; ROSA26^mTmG/+^* mice at postnatal (P) day 0. No difference in saccule formation was noted between control and mutant lungs. (D) Immunostaining of lung sections from control and *Miro^f/f^; Sox9^Cre/+^; ROSA26^mTmG/+^* mice at P5. Specification of lung cell types were unaffected.

**Figure 5–figure supplement 1.**
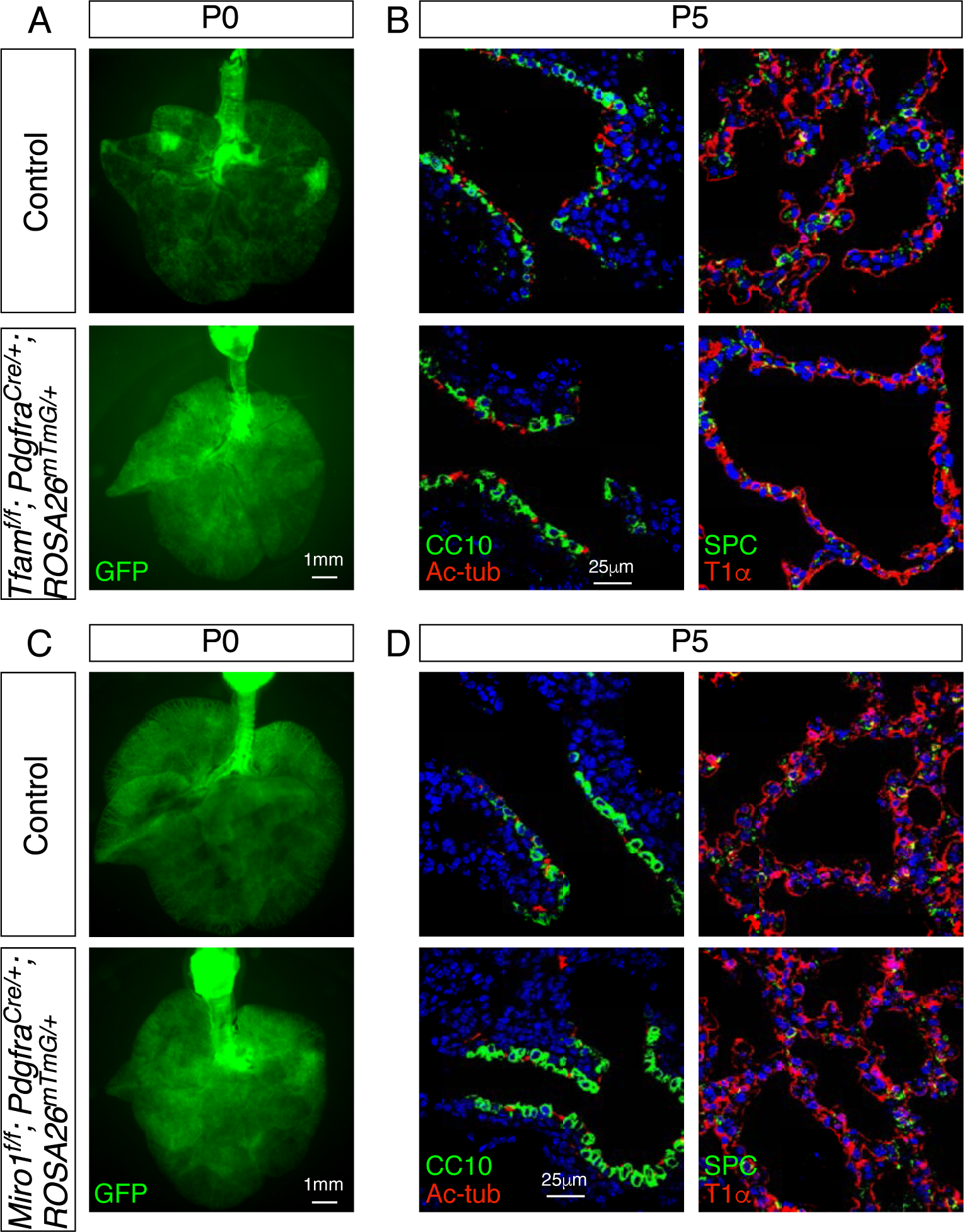
Removal of *Tfam* or *Miro1* in mouse lung fibroblasts/myofibroblasts does not perturb saccule formation or cell type specification. (A) Surface view of dissected lungs from *Pdgfra^Cre/+^; ROSA26^mTmG/+^* (control) and *Tfam^f/f^; Pdgfra^Cre/+^; ROSA26^mTmG/+^* mice at postnatal (P) day 0. No difference in saccule formation was noted between control and mutant lungs. (B) Immunostaining of lung sections from control and *Tfam^f/f^; Pdgfra^Cre/+^; ROSA26^mTmG/+^* mice at P5. Specification of lung cell types were unaffected. (C) Surface view of dissected lungs from control and *Miro1f/f; Pdgfra^Cre/+^; ROSA26^mTmG/+^* mice at postnatal (P) day 0. No difference in saccule formation was noted between control and mutant lungs. (D) Immunostaining of lung sections from control and *Miro^f/f^; Pdgfra^Cre/+^; ROSA26^mTmG/+^* mice at P5. Specification of lung cell types were unaffected.

**Figure 5–figure supplement 2.**
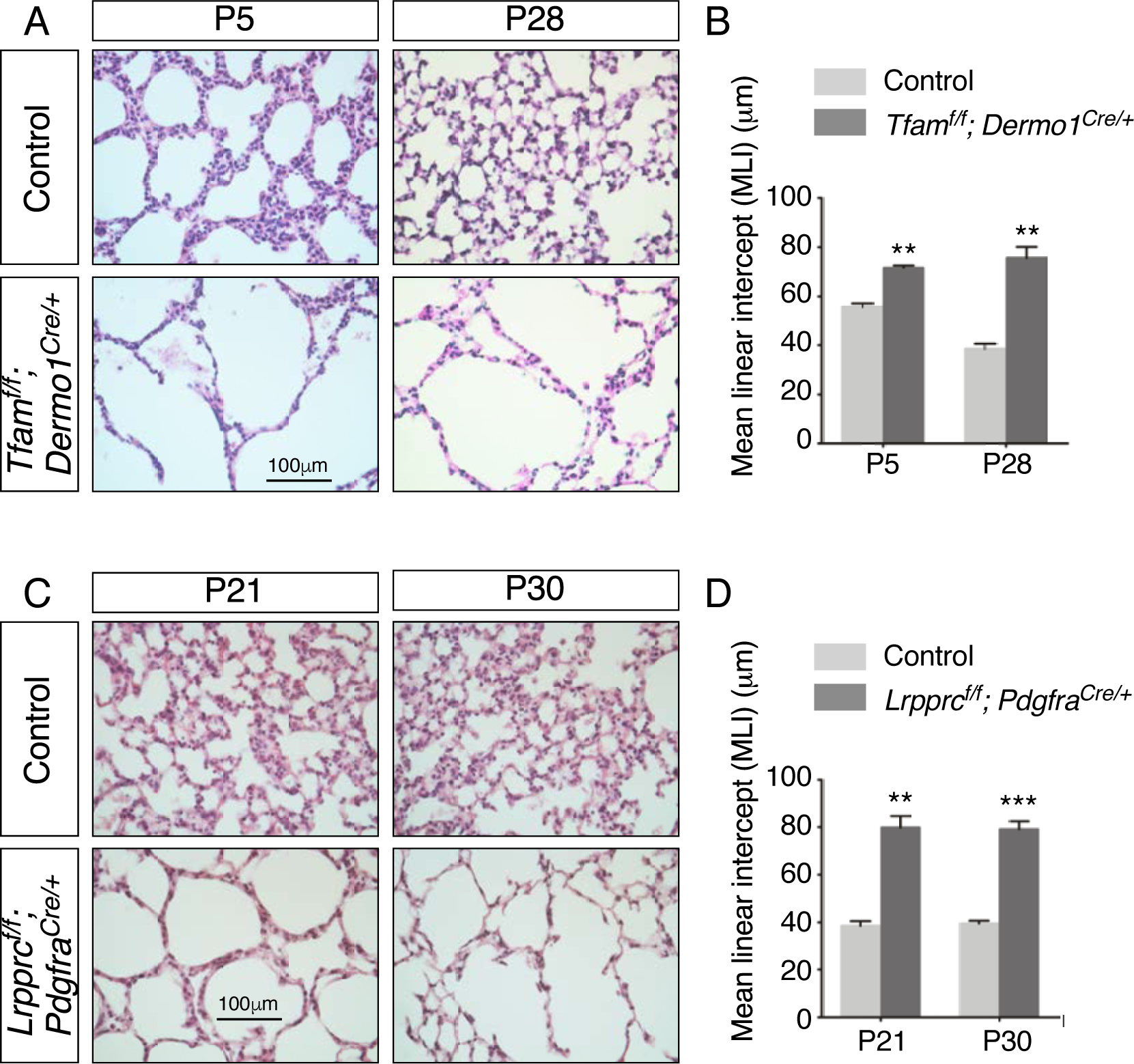
Inactivation of *Tfam* or *Lrpprc* in mouse lung fibroblasts/myofibroblasts leads to alveolar defects. (A) Hematoxylin and eosin-stained lung sections of control and *Tfam^f/f^; Dermo1^Cre/+^* mice at different postnatal (P) stages as indicated. Histological analysis revealed the presence of enlarged saccules and defective development of secondary septa in the mutant lungs. (B) Measurement of the MLI in control and *Tfam^f/f^; Dermo1^Cre/+^* lungs at P5 and P28 (n = 3 for each group). The MLI was increased in *Tfam*-deficient lungs. (C) Hematoxylin and eosin-stained lung sections of control and *Lrpprc^f/f^; Pdgfra^Cre/+^* mice at P21 and P30. Saccules were enlarged with loss of secondary septation in the mutant lungs. (D) The MLI was increased in *Lrpprc*-deficient lungs at P21 and P30 (n = 3 for each group). All values are mean SEM. (*) p<0.05; (**) p<0.01; (***) p<0.001; ns, not significant (unpaired Student’s t-test).

